# A single-cell reference map for human blood and tissue T cell activation reveals functional states in health and disease

**DOI:** 10.1101/555557

**Authors:** Peter A. Szabo, Hanna Mendes Levitin, Michelle Miron, Mark E. Snyder, Takashi Senda, Jinzhou Yuan, Yim Ling Cheng, Erin C. Bush, Pranay Dogra, Puspa Thapa, Donna L. Farber, Peter A. Sims

## Abstract

Human T cells coordinate adaptive immunity by localization in diverse tissue sites, though blood T cells are the most readily studied. Here, we used single-cell RNA-seq to define the functional responses of T cells isolated from human lungs, lymph nodes, bone marrow, and blood to TCR-stimulation. We reveal how human T cells in tissues relate to those in blood, and define activation states for CD4^+^ and CD8^+^T cells across all sites, including an interferon-response state for CD4^+^T cells and distinct effector states for CD8^+^T cells. We further show how profiles of individual tumor-associated T cells can be projected onto this healthy reference map, revealing their functional state.

## INTRODUCTION

T lymphocytes coordinate adaptive responses and are essential for establishing protective immunity and maintaining immune homeostasis. Activation of naïve T cells through the antigen-specific T cell receptor (TCR) initiates transcriptional programs that drive differentiation of lineage-specific effector functions; CD4^+^T cells secrete cytokines to recruit and activate other immune cells while CD8^+^T cells acquire cytotoxic functions to directly kill infected or tumor cells. Most of these effector cells are short-lived, although some develop into long-lived memory T cells which persist as circulating central (TCM) and effector-memory (TEM) subsets, and non-circulating tissue resident memory T cells (TRM) in diverse lymphoid and non-lymphoid sites^1–4^. Recent studies in mouse models have established an important role for CD4^+^ and CD8^+^TRM in mediating protective immunity to diverse pathogens^2,5–7^. Defining how tissue site impacts T cell function is therefore important for targeting T cell immunity.

In humans, most of our knowledge of T cell activation and function derives from the sampling of peripheral blood. Recent studies in human tissues have revealed that the majority of human T cells are localized in lymphoid mucosal and barrier tissues^8^ and that T cell subset composition is a function of the specific tissue site^9,10^. Human TRM cells can be defined based on their phenotypic homology to mouse TRM and are distinguished from circulating T cells in blood and tissues by a core transcriptional and protein signature^10–13^. However, the role of tissue site in determining T cell functional responses, and a deeper understanding of the relationship between blood and tissue T cells beyond composition differences are key unanswered questions in human immunology.

The functional responses of T cells following antigen or pathogen exposure have been largely defined in mouse models, and are generally classified based on whether or not they secrete specific cytokines or effector molecules. Effector CD4 T cells comprise different functional subtypes (Th1 cells secrete IFN-γ and IL-2; Th2 secrete IL-4, 13; Th17 secrete IL-17, etc.)^14^, while effector CD8 T cells secrete pro-inflammatory cytokines (IFN-γ,TNF-α) and/or cytotoxic mediators (perforin and granzymes)^15^. Certain conditions can lead to inhibition of functional responses; for example, CD4^+^T cells encountering self-antigen become anergic and fail to produce IL-2, while CD8^+^T cells responding to chronic infection, tumors, or lacking CD4^+^T cell help become functionally exhausted, and express multiple inhibitory molecules (e.g., PD-1, LAG3)^16–18^. While human T cells can produce similar cytokines, effector and inhibitory molecules as mouse counterparts^19–22^, the full complement of functional responses for human T cells in tissues has not been elucidated. Establishing a comprehensive baseline of healthy T cell states in humans is essential for defining dysregulated and pathological functions of T cells in disease.

Single cell transcriptome profiling (scRNA-seq) has enabled high resolution mapping of cellular heterogeneity, development, and activation states in diverse systems^23,24^. This approach has been applied to analyze human T cells in diseased tissues^25,26^ and in response to immunotherapies in cancer ^27^; however, the baseline functional profiles of human T cells in healthy blood and tissues have not been defined. We have established a tissue resource where we obtain multiple lymphoid, mucosal, and other peripheral tissue sites from human organ donors^9–11,13,28,29^, enabling study of T cells across different anatomical spaces. Here, we used scRNA-seq of over 50,000 resting and activated T cells from lung (LG), lymph nodes (LN), bone marrow (BM) and blood, along with innovative computational analysis to define cellular states of homeostasis and activation of human blood and tissue-derived T cells. We reveal how human T cells in tissues relate to those in blood, and identify a conserved tissue signature and activation states for human CD4^+^ and CD8^+^T cells conserved across all sites. We further show how scRNA-seq profiles of T cells associated with human tumors can be projected onto this healthy baseline dataset, revealing their functional state. Our results establish a comprehensive high dimensional dataset of human T cell homeostasis and function in multiple sites, from which to define the origin, composition and function of T cells in disease.

## RESULTS

### High Resolution analysis of human T cells in tissues and comparison to blood

We obtained BM, LN, and LG as representative primary lymphoid, secondary lymphoid and mucosal tissue sites, respectively, from two adult organ donors who met the criteria of health for donation of physiologically healthy tissues for lifesaving transplantation, being free of chronic disease, cancer, and infections (Supplementary Table 1). For comparison, we obtained blood from two healthy adult volunteers. CD3^+^T cells isolated from tissues and blood were cultured in media alone (“resting”) or in the presence of anti-CD3/anti-CD28 antibodies (“activated”) (Fig. 1a). Single cells were encapsulated for cDNA synthesis and barcoding using the 10x Genomics Chromium system, followed by library construction, sequencing, and computational identification of T cells (Supplementary Fig. 1, Supplementary Tables 2,3).

**Figure 1:**
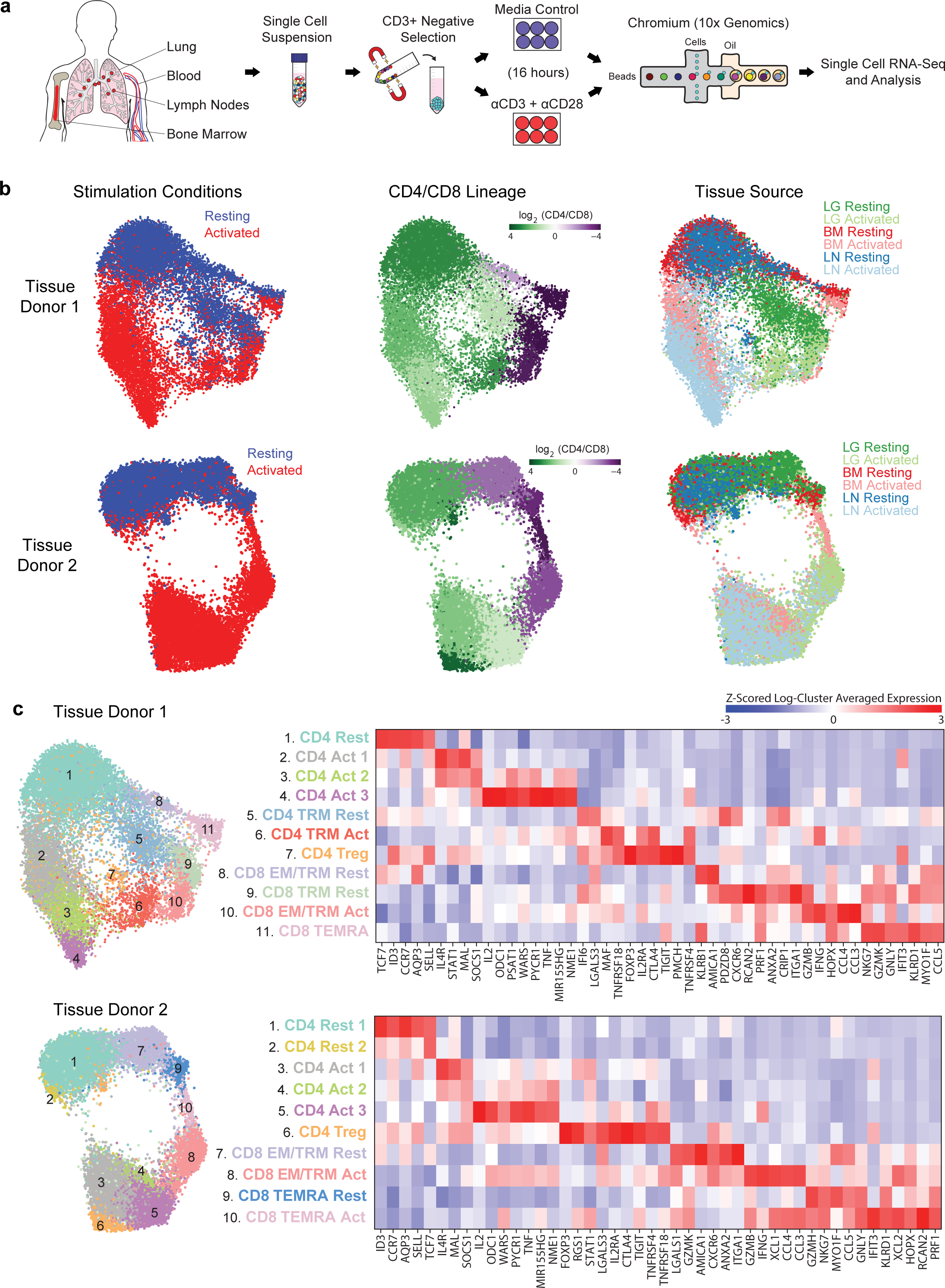
Single-cell RNA-seq analysis of resting and activated T cells from multiple tissue sites in each individual donor. (a) Experimental workflow for single-cell analysis of T cells from human tissues and blood including magnetic negative selection of CD3^+^ cells, *in vitro* culture and activation, and Chromium 3’-scRNA-seq. (b) UMAP embeddings of merged scRNA-seq profiles from resting and activated T cells from lung (LG), bone marrow (BM), and lung-draining lymph node (LN) in each of two organ donors colored by resting/activated condition, CD4/CD8 expression ratio (all cells in a given cluster assigned the same average value), and tissue source. (c) Identification of T cell subpopulations. UMAP embeddings colored by expression cluster along with heatmaps showing the z-scored average expression of differentially expressed marker genes for each cluster. Subsets designated based on resting (“rest”) or activated (“act”) condition and expression of known markers denoting effector memory (TEM), tissue resident memory (TRM), terminally differentiated effector cells (TEMRA), and regulatory T cells (Treg).

We initially analyzed tissue T cell populations from the two individual donors, comprising six samples per donor (resting and activated samples from three tissue sites). We merged all data for each donor, performed unsupervised community detection^30^ to cluster the data based on highly-variable genes (Supplementary Table 4), and projected cells in two dimensions using Uniform Manifold Approximation and Projection (UMAP)^31^. For both donors, the dominant sources of variation between cells were activation state (vertical axis) and CD4/CD8 lineage (horizontal axis) (Fig. 1b). Tissue site was also a source of variability; T cells from BM and LN co-clustered while LG T cells were more distinct (Fig. 1b), consistent with T cell subset composition differences in these sites from phenotype analysis (Supplementary Figure 2 and previous studies^10,13,32^).

Differential gene expression from the scRNA-seq data resolved T cell subsets and functional states within and between sites and lineages into 10-11 clusters (Fig. 1c, Supplementary Tables 5,6). CD4^+^T cells comprised 6-7 clusters: resting cells expressing *CCR7*, *SELL* and *TCF7*, (corresponding to naïve or TCM cells); three activation-associated clusters expressing *IL2*, *TNF*, and *IL4R* at different levels; TRM-like resting and activated clusters expressing canonical TRM markers *CXCR6* and *ITGA1^13,33^*; and a distinct regulatory T cell (Treg) cluster expressing Treg-defining genes *FOXP3*, *IL2RA*, and *CTLA4* (Fig. 1c). CD8^+^T cells comprised four clusters distinct from CD4^+^T cells and included: two TEM/TRM-like clusters expressing *CCL5*, cytotoxicity-associated genes (*GZMB, GZMK*), and TRM markers (*CXCR6*, *ITGA1*); an activated TRM/TEM cluster expressing *IFNG, CCL4, CCL3*; and clusters representing terminally differentiated effector cells (TEMRA) expressing cytotoxic markers *PRF1* and *NKG7* (Fig. 1c). In terms of tissue distribution, TRM cells were largely in the lung, Tregs were primarily identified in LN, while TEMRA cells were enriched in BM (consistent with phenotype analysis, Supplementary Fig. 2); the remaining resting and activated CD4^+^ and CD8^+^ clusters derived from all sites (Fig. 1b,c). These results show subset-specific profiles in human tissues, but suggest similar ^13^activation profiles across sites.

To assess how blood T cells relate to those in tissue, we performed scRNA-seq analysis of resting and activated blood T cells from two adult donors, and projected the merged data onto the UMAP embeddings of T cells from each tissue donor (Fig. 2a,b, Online Methods). The majority of blood T cells co-localized with resting or activated T cells from BM but did not exhibit substantial overlap with LG or LN T cells from either donor, particularly in the resting state (Fig. 2a,b). We also quantified the number of blood T cells that were transcriptionally similar to CD4^+^ and CD8^+^T cells from each tissue within resting or activated samples (Fig. 2c,d, Online Methods). Resting blood T cells were highly represented among CD4^+^ and CD8^+^T cells in BM (Fig. 2c, d). Interestingly, a substantial number of unstimulated blood T cells projected onto activated CD4^+^T cells in BM for both donors (Fig. 2c,d, left panels). In contrast, activated blood T cells were strongly represented among activated CD4^+^T cells for all tissue sites and in LN for CD8^+^T cells (Fig. 2c,d; right panels). Consistent results were obtained analyzing each blood sample separately (Supplementary Fig. 3). These results suggest that blood T cells are fundamentally distinct from tissue T cells and may persist in a more activated basal state than in tissues, while activated blood and tissue-derived T cells share common signatures.

**Figure 2:**
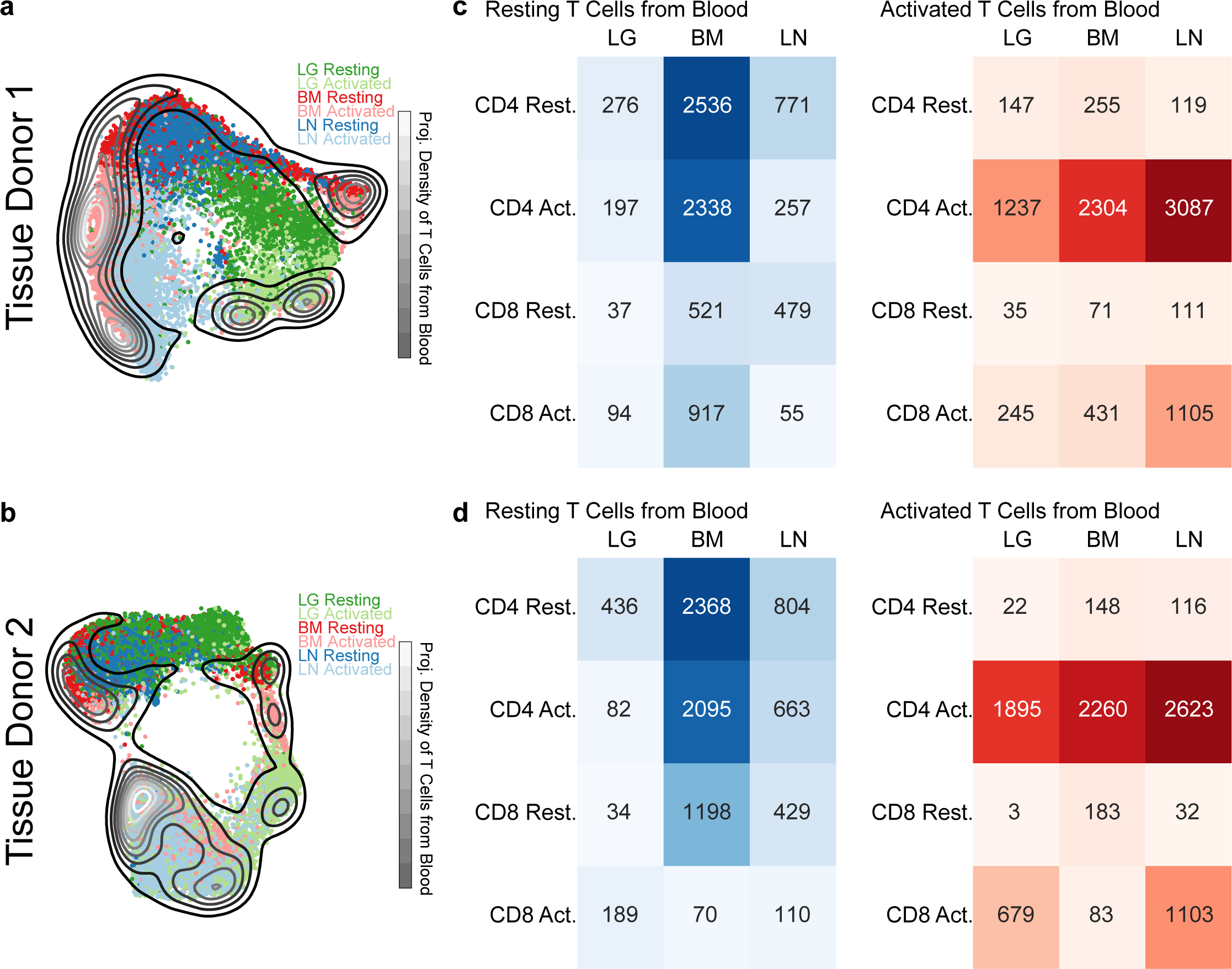
Comparison of blood and tissue T cells. (a) UMAP embedding of T cells from tissue donor 1 colored by tissue and overlaid with a contour plot corresponding to the UMAP projection of the combined resting and activated T cells from two blood donors onto the tissue embedding. (b) Same as (a) for organ donor 2. (c) Heatmaps showing the number of blood T cells that project most closely to each tissue/stimulation status combination in the tissue donor 1 UMAP embedding. (d) Same as (c) for tissue donor 2.

**Figure 3:**
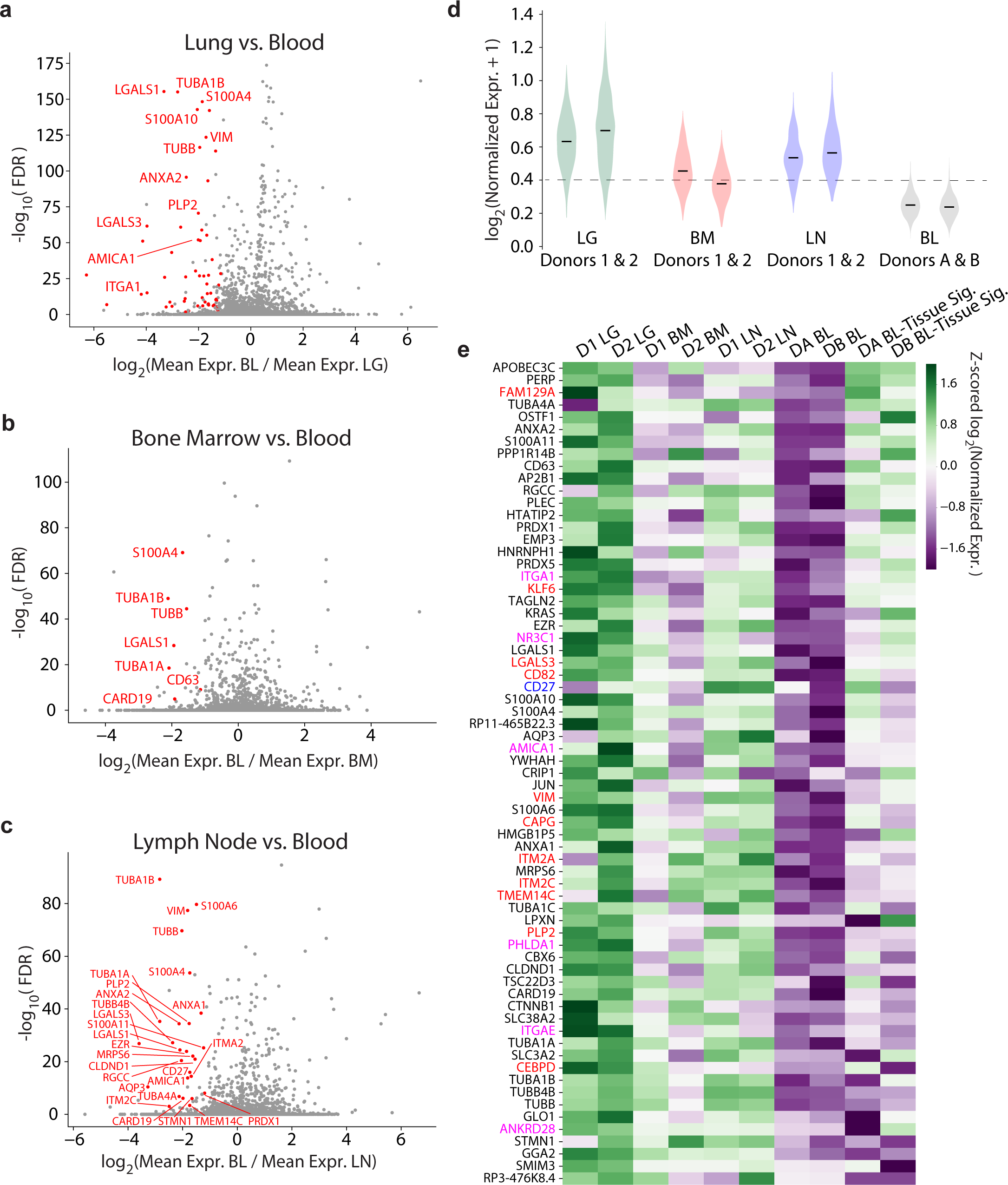
Identification of a tissue gene signature for resting memory T cells. (a) Volcano plot showing the average log-fold-change and average Benjamini-Hochberg-corrected p-values (FDR) for pairwise differential expression between CCL5^+^ T cells from each resting LG sample and each resting blood sample. Genes with negative log-fold-change are more highly expressed among CCL5^+^ cells in LG, with several highlighted in red. (b) Same as (a) for comparison of resting CCL5^+^ T cells in BM and blood. (c) Same as (a) for comparison of resting CCL5^+^ T cells in LN in blood. (d) Violin plot showing the distributions of the average expression of all genes with two-fold higher expression (on average) in any tissue compared to blood and average FDR < 0.05 in any tissue for the resting CCL5^+^ T cells in each tissue and blood sample. The dashed line marks one standard deviation below the mean for average expression of this signature for all tissues (note a small number of blood cells fall above this line). (e) Heatmap shows z-scored average expression for all genes in the tissue signature from (d) among the resting CCL5^+^ T cells from each tissue and blood sample plus that of the rare blood subpopulation from (d), which expresses high levels of certain genes. Previously identified TRM-associated genes from bulk RNA-seq studies are highlighted in red (enriched in CD69^+^ vs. CD69^−^), blue (enriched in CD69^+^/CD103^+^ vs. CD69^+^/CD103^−^), and magenta (enriched in both CD69^+^ vs. CD69^−^ and CD69^+^/CD103^+^ vs. CD69^+^/CD103^−^) (See online methods).

### A universal gene signature distinguishes tissue T cells from blood

The major transcriptional differences between tissue and blood T cells based on population RNAseq originate from the presence of TRM in tissues^13^. Because scRNA-seq enables high resolution detection of gene expression differences that can be unambiguously traced to specific T cells, we investigated whether there were intrinsic features of tissue T cells that distinguished them from blood. Resting memory T cells in tissues and blood express high levels of CCL5 (Supplementary Fig. 4, Online Methods), a marker of CD8^+^TEM cells^34^, enabling direct comparison of gene expression between similar subsets. We identified a similar complement of genes that were highly expressed in TEM cells from each tissue compared to blood (Fig. 3a-c). Interestingly, these tissue-intrinsic genes include those associated with microtubules and the cytoskeleton (tubulin-encoding genes T*UBA1A, TUBA1B, TUBB, TUBB4B; S100A4*) and genes encoding cell matrix, membrane scaffolding, and adhesion molecules (*VIM* or vimentin, galectins *LGALS1/LGALS3*, *AMICA1, ITM2C, EZR,* annexins *ANXA1/ANXA2*) (Fig. 3a-c). TRM signature genes including ITGA1 and ITGAE were also upregulated in tissues compared to blood, particularly in the lung (Fig. 3a-c). These findings suggest that localization of T cells in tissues likely involves structural changes in the cell that facilitate interactions with tissue matrix.

We compared the single-cell distribution of average expression of tissue signature genes in the blood and three tissues (Fig. 3d). CCL5^+^TEM cells from all three tissues (both donors) express higher levels of tissue signature genes compared to blood, though LG and LN T cells have higher expression than those from BM (Fig, 3d). Notably, a minute fraction of blood TEM cells (<0.5%) express this tissue signature at levels comparable to that in LN (within one standard deviation of the mean for all tissues). Shown in a heat map are the relative expression levels for genes within the tissue signature including TRM signature genes^13^ cytoskeletal, cell-matrix interactions, cell division, apoptotic, and signaling genes (Fig. 3e). Gene expression is highest in LG followed by LN and BM expressing only a subset of tissue-associated genes; the outlier subpopulation from blood expresses a fraction (<40%) of tissue signature genes at levels comparable to those in tissues (Fig. 3e). Together, these results show that tissue T cells express genes associated with infiltration and localization in tissues along with residency markers, while blood contains only trace numbers of cells expressing these genes.

### Defining functional states common to blood and tissue T cells

The clustering analysis above suggested that activated T cells were more similar across sites than resting counterparts. To uncover gene expression patterns that were conserved across T cell populations in different tissues, we applied a new analytical method called single-cell Hierarchical Poisson Factorization (scHPF)^35^. The scHPF algorithm identifies a small number of expression patterns, called factors that vary coherently across cells. These factors can represent discrete, subpopulation-specific or continuous programs like T cell activation that are expressed as a gradient across cells in different stages of a biological process. We applied scHPF to merged resting and activated T cells from each tissue and donor separately and hierarchically clustered the resulting factors (Online Methods, Fig. 4a, Extended Data Fig. 4a, Supplementary Fig. 5). This analysis revealed seven gene expression modules (3 resting and 4 activated/functional) that were highly conserved across tissues and donors, for which the highest scoring genes formed interpretable gene signatures (Fig. 4a, Extended Fig. 4a, Supplementary Table 7). The three modules associated with a resting state (Fig. 4a) included a Treg module defined by canonical genes (*FOXP3*, *CTLA4*, *IRF4*, *TNFRSF4* (OX40)^36^); a putative resting CD4^+^ Naïve/Central memory (NV/CM) module enriched in CD4^+^T cells and defined by genes associated with lymphoid homing, egress and quiescence (*SELL*, *KLF2*, *LEF1,* respectively), while the CD4^+^/CD8^+^ Resting module was distinguished by expression of *IL7R*, a receptor required for T cell survival^37,38^, and *AQP3*, which encodes a water channel protein of unclear function in lymphocytes^39^. Importantly, the CD4^+^/CD8^+^ Resting module did not contain factors from blood and had the highest enrichment for the tissue signature identified in Fig. 3 (Supplementary Fig. 6).

**Figure 4:**
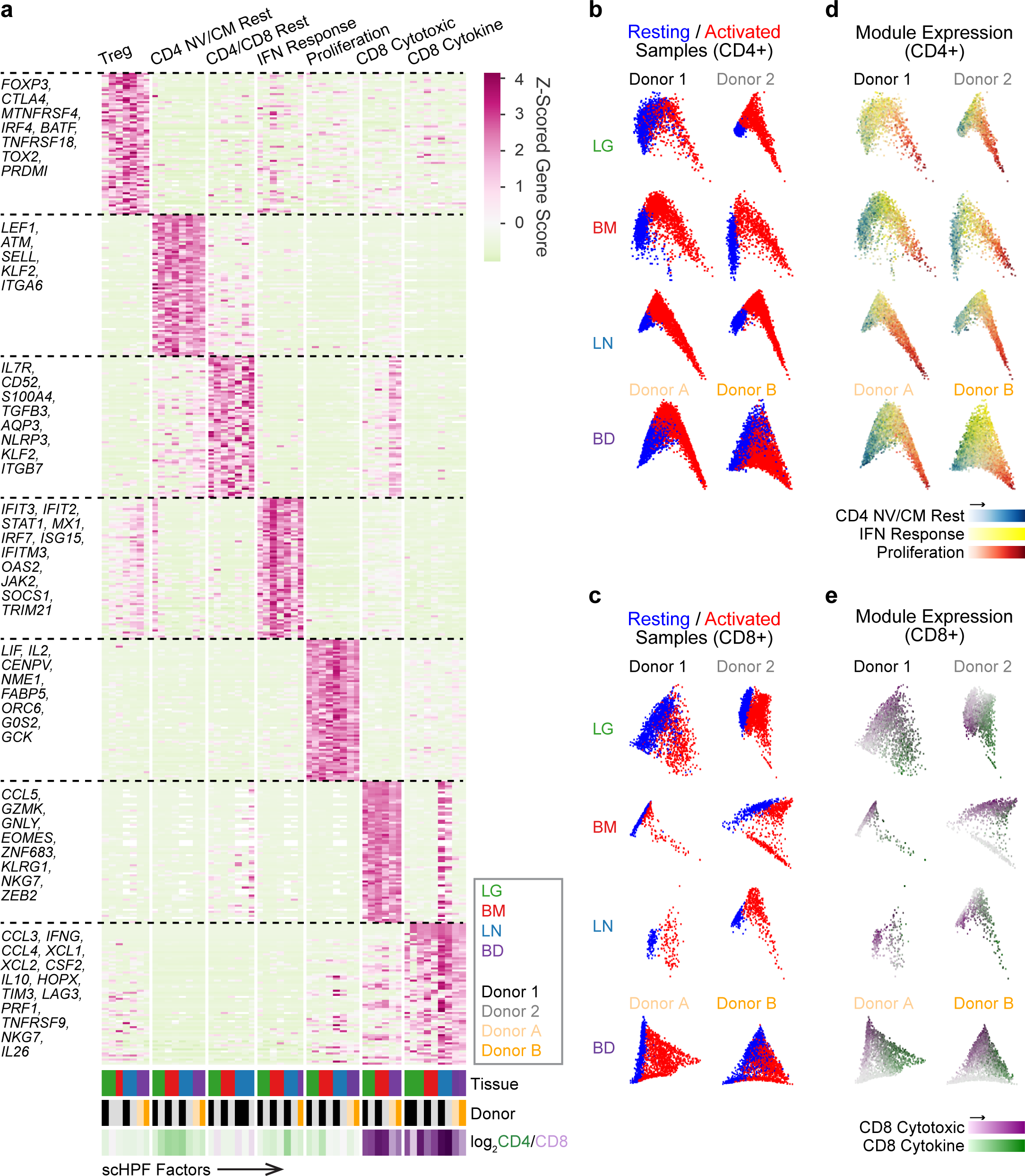
Defining conserved transcriptional states in resting and activated T cells by single-cell Hierarchical Poisson Factorization (scHPF). (a) Heatmap shows gene scores for the top genes (rows) in each expression module identified by clustering scHPF factors (columns) that were computed in separate analyses of cells from each tissue and donor (Extended Figure 4a, Online Methods). Selected genes are indicated to the left, and complete lists of top genes are available in Supplementary Table 7. Color bars at the bottom of the heatmap indicate each factors’ tissue of origin, donor of origin, and CD4/CD8 bias. (NV/CM=naïve or TCM). (b) Diffusion maps of CD4^+^ T cells in each tissue and donor, with cells colored by sample origin as resting (blue) or activated (red). (c) Same as (b) but for CD8+T cells. (d) Diffusion maps of CD4^+^T cells from (b), with cells colored by their average expression of the top genes from scHPF expression modules. Colors for different modules (CD4 NV/CM Resting, IFN Response, Proliferation) were blended using the RGB color model. (e) Diffusion maps of CD8^+^T cells from (c), with cells colored by their average expression of the top genes from scHPF expression modules (CD8 Cytotoxic and CD8 Cytokine).

There were four modules associated with T cell activation and/or function, some of which were lineage-specific. A Proliferation module expressed by activated CD4^+^ and CD8^+^ lineages included genes associated with T cell activation/proliferation (*IL2, LIF*) and cell division (*CENPV*, *G0S2*, *ORC6*) (Fig. 4a). This module was also marked by expression of *NME1*, a metastasis suppressor/endonuclease-encoding gene^40^ not previously associated with T cells (Fig. 4a). An Interferon (IFN) Response module enriched among activated CD4^+^T cells included multiple gene families associated with canonical IFN responses^41–43^ (*IFIT3*, *IFIT2*, *STAT1*, *MX1*, *IRF7*, and *JAK2*). In contrast, CD8^+^T cell-enriched modules included a Cytotoxic module containing genes associated with cytotoxicity (*GNLY*, *GZMK*) and transcription factors associated with effector/memory differentiation (*ZEB2*, *EOMES, ZNF683)*^43–45^, and a Cytokine module with genes encoding chemokines and cytokines (*CCL3*, *CCL4*, *CCL20*, *IFNG*, *IL10*, *TNF*), inhibitory molecules (*LAG3*, *CD226* (TIGIT), *HAVCR2* (TIM3)), and the widely expressed homeobox protein *HOPX* ^46^. These results indicate a limited spectrum of functional states for human T cells across blood and tissue sites.

To understand how these gene modules correspond to resting and activated states in CD4^+^ and CD8^+^T cells, we visualized the average expression of their top-ranked genes on diffusion maps for each donor and tissue (Fig. 4b-e, Online Methods). This visualization defined activation trajectories with resting T cells on the left (blue) and activated T cells projecting to the right (red, Fig. 4b,c). In all four sites and individuals, module expression for CD4^+^T cell was positioned along activation trajectories from CD4 NV/CM Resting (left) to IFN-Response (middle) to Proliferation (right) (Fig. 4d). Expression of genes within the Proliferation module co-localized with peak expression of *NME1* and *IL2RA* (Extended Data Fig. 4b,c), while the IFN Response module genes exhibited peak expression at the middle of the trajectory as exemplified by *IFIT3* expression (top ranked gene) (Extended Data Fig. 4d), suggesting a potential intermediate activation state. In CD8^+^T cells, the Cytokine module localized in the most activated cells for all sites also shown by *IFNG* expression (Fig. 4e, extended data Fig. 4e), while the Cytotoxic module was expressed among resting and activated cells (Fig. 4e). Therefore, scHPF takes an unbiased approach to uncover major functional states, reference signatures and activation trajectories for human T cells that are conserved across sites.

### CD4^+^ T cell activation states result from distinct responses to TCR and type II IFN signaling

The functional states identified for human CD8^+^T cells in Fig. 4 were consistent in with those seen *in vivo* in mouse infection models^15^. By contrast, the modules identified for CD4^+^T cell activation revealed markers and functional states not typically associated with effector CD4^+^T cells. We therefore assessed expression kinetics of the top-scoring genes in the Proliferation and IFN Response modules, *NME1* and *IFIT3*, respectively, during the course of T cell activation *ex vivo* by qPCR. Expression of *NME1* transcripts rapidly increased after TCR-stimulation, peaking between 16-24hrs and remaining elevated for up to 72hrs, for both CD4^+^ and CD8^+^T cells compared to unstimulated controls, a pattern of expression similar to the canonical T cell activation marker *IL2RA* (Fig. 5a). Notably, the extent of activation-associated upregulation of *NME1* transcripts was greater in CD4^+^ compared to CD8^+^T cells, while *IL2RA* was more upregulated in CD8^+^T cells (Fig. 5a). At the protein level, NME1 expression increased in CD4^+^ and CD8^+^T cells after TCR-mediated stimulation from 24-120 hours (Fig. 5b, upper), and with each successive round of T cell proliferation, while CD25 was expressed similarly independent of cell division (Fig. 5b, lower). These results establish NME1 expression as a marker of T cell activation, coupled to the extent of proliferation.

**Figure 5:**
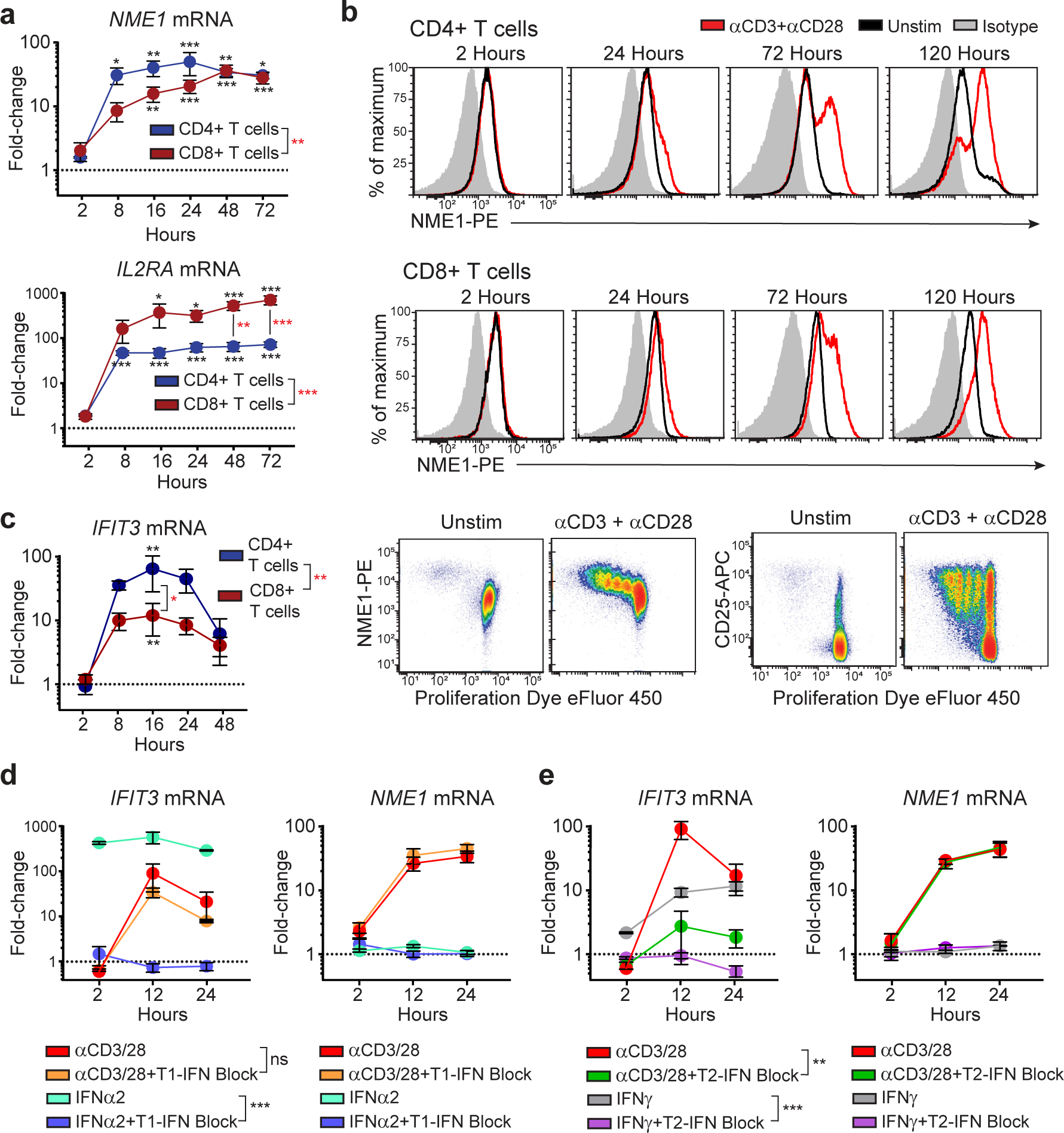
NME1 and IFIT3 induction during CD4^+^T cell activation and the role of IFN-signaling. (a) Expression of *NME1* and *IL2RA* mRNA by blood CD4^+^ or CD8^+^T cells after stimulation with anti-CD3/anti-CD28 antibodies by qPCR. Data shown as mean fold-change (±SEM) relative to unstimulated CD4^+^ or CD8^+^T cell controls (dotted line) from 4 individuals (independent experiments). Statistical analysis between stimulated and unstimulated cells (black *) or CD4^+^ and CD8^+^T cells (red *) made by two-way ANOVA with Sidak test for multiple comparisons. (b) Intracellular NME1 protein expression by blood T cells after stimulation for indicated timepoints (red) compared to unstimulated (black) and isotype control (gray). Bottom row: CD25 and NME1 expression by proliferating CD3^+^T cells after 5 days of stimulation. Data are representative of 4 individuals. (c) Expression of *IFIT3* mRNA in blood T cells by qPCR after TCR-stimulation, shown as mean fold-change (±SEM) relative to unstimulated controls (dotted line) for four individuals. Two-way ANOVA with Sidak test for multiple comparisons was used for statistical comparisons (black *, stimulated versus unstimulated) or (red *, CD4^+^ versus CD8^+^T cells). (d) *IFIT3* or *NME1* mRNA expression in CD4^+^T cells after culture with anti-CD3/anti-CD28 or IFNα2 (1000 units/mL) +/-type I IFN neutralizing antibody cocktail or (e) IFNγ (10 ng/mL) +/-anti-IFNγ/anti-IFNγR1 antibodies (1 ug/mL each), shown as mean fold-change (±SEM) relative to unstimulated controls (dotted line) for 3 individuals. Statistical comparisons made by two-way ANOVA. For all panels: “ns” denotes not significant; * *p ≤* 0.05; ** *p ≤* 0.01; *** *p ≤* 0.001.

In contrast to *NME1/IL2RA* upregulation, expression of *IFIT3* transcripts showed biased and transient upregulation by CD4^+^T cells following TCR-stimulation, peaking at 16hrs and returning to near baseline levels by 48hrs post-stimulation (Fig. 5c). As *IFIT3* expression is associated with responses to IFN^41^, we assessed the kinetics of *IFIT3* upregulation after IFN compared to TCR-mediated signaling and the contribution type I or type II IFN signaling to TCR-triggered *IFIT3* induction. In response to IFN-α (type I) or IFN-γ (type II), *IFIT3* was more rapidly (within 2hrs) and persistently upregulated compared to TCR stimulation (Fig. 5d,e). Inhibiting type I IFN signaling (using neutralizing antibodies for type I IFNs and IFNαR2) abrogated IFN-αinduced *IFIT3* upregulation, as expected, but did not affect TCR-mediated upregulation of *IFIT3* by CD4^+^T cells (Fig. 5d). However, blockade of type II IFN signaling via a combination of anti-IFNγ and anti-IFNγR1 antibodies inhibited upregulation of *IFIT3* induced by TCR-mediated activation, as well as that induced by culture with IFN-γ (Fig. 5e). Notably, blocking type II (or type I) IFN signaling did not inhibit T cell activation as assessed by induction of *NME1* transcript expression, and addition of IFN-α or –γ did not induce *NME1* expression (Fig. 5d,e). These results establish that the IFN-responsive state suggested by the scRNA-seq trajectories is recapitulated in real-time as part of an intermediate activation state driven by TCR-triggered IFN-γproduction.

### Defining T cell activation states in cancer through projecting on the high resolution reference map

Although there have been several large-scale scRNA-seq studies of disease-associated T cells, these data are generally not placed in the context of T cell activation in healthy individuals. To demonstrate the utility of our resource as a reference point for human disease, we used UMAP to project recently reported scRNA-seq profiles of tumor-associated T cells from four different human cancers onto our map of T cell activation states. Fig. 6a shows a merged UMAP embedding of our entire data set colored by tissue site, donor, stimulation, cluster-level CD4/CD8 status, and CCL5 expression, indicative of effector status. We projected scRNA-seq profiles of tumor-associated T cells from four different human cancers^27,47–49^ (non-small cell lung cancer (NSCLC), colorectal cancer (CRC), breast cancer (BC), and melanoma (MEL)) onto this embedding to compare each tumor-associated T cell to healthy T cells (Fig. 6b,c). We also investigated expression of activation state and lineage markers in the healthy T cell embedding and tumor projections (Fig. 6c). Tumor-associated CD8^+^T cells project onto healthy CD8^+^T cells from all sites in both resting and activated states (Fig. 6b). Moreover, genes associated with TRM (CXCR6) and the Cytotoxic and Cytokine modules are all represented among tumor-associated CD8^+^T cells (Fig. 6c, Extended data fig. 6, Supplementary Fig. 7,8). By contrast, tumor-associated CD4^+^T cells projected mostly onto resting blood and tissue T cells (Fig. 6b), while CD4^+^T cell activation states and associated markers (*NME1*, IFIT3) were largely absent (Fig. 6c). This analysis reveals that tumor-associated T cells contain activated CD8^+^T cells, but lack the presence of functionally activated CD4 T cells.

**Figure 6:**
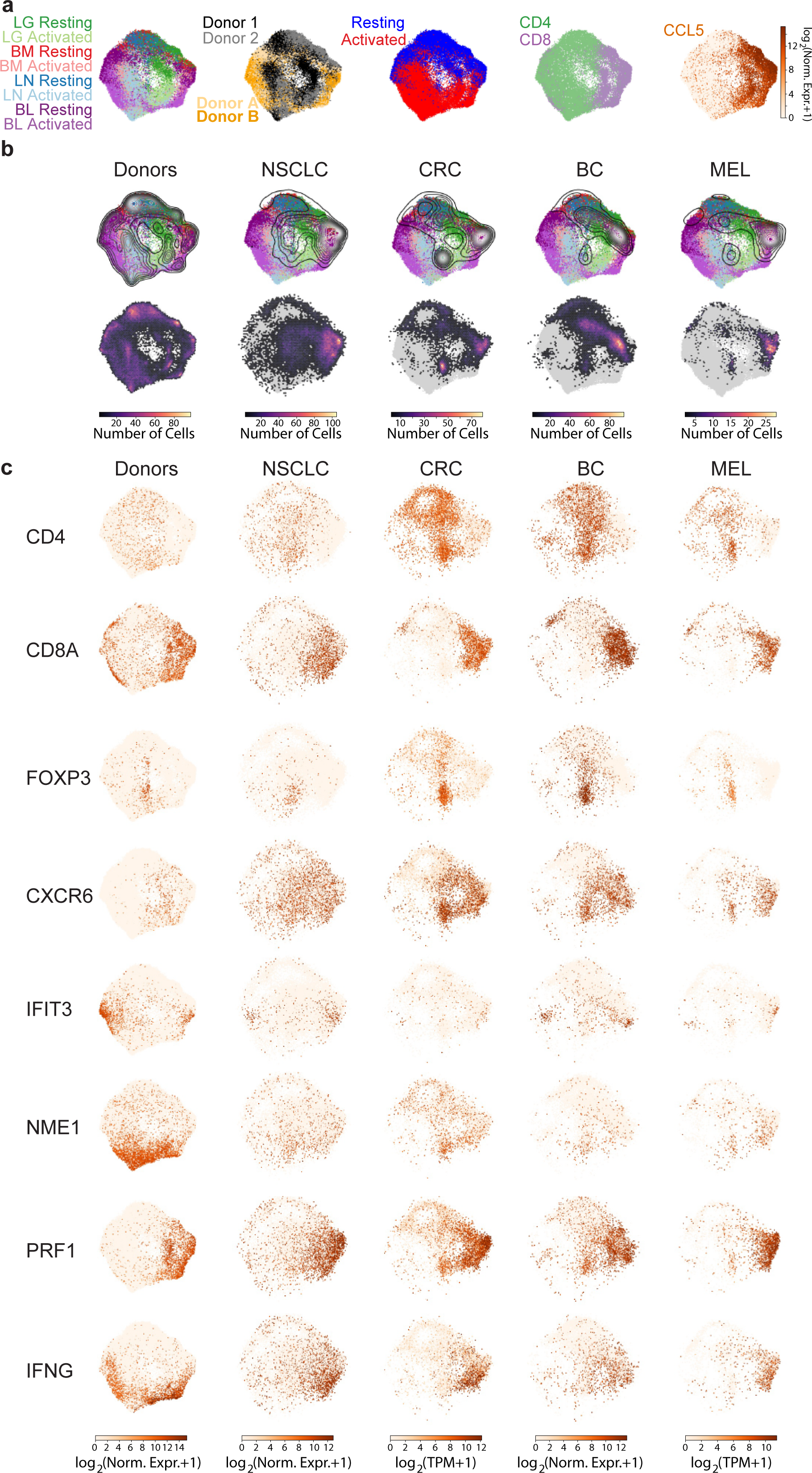
Comparison of tumor-associated T cells to the reference map of healthy human T cell activation. (a) Merged UMAP embedding for the entire healthy T cell scRNAseq dataset including resting and activated tissue T cells (two donors) and blood T cells (two individual) colored by sample source, donor, resting/activated condition, CD4/CD8 status (CD4-enriched, green; CD8-enriched, purple), and CCL5 expression indicating TEM cells. (b) Merged UMAP embedding for the entire dataset overlaid with contour plots indicating kernel density estimates for the projection of T cells derived from organ/blood donors (column 1), non-small cell lung cancer (NSCLC) tissue (column 2), colorectal cancer (CRC) tissue (column 3), breast cancer (BC) tissue (column 4), and melanoma (MEL) tissue (column 5). Note that these probability densities can be compared within each projection, but cannot be quantitatively compared across projections. (c) Same as b) but overlaid with a two-dimensional hexbin histogram for each projection. Histograms have been normalized to account for differences in cell numbers across datasets and therefore can be compared quantitatively across projections. (d) Individual cells in the UMAP embedding (column 1) for the entire healthy T cell dataset and UMAP projections (columns 2-5) for NSCLC, CRC, BC, and MEL tissue T cells colored by expression of *CD4*, *CD8A*, *FOXP3* (Treg marker), *CXCR6* (TRM marker), *IFIT3* (IFN response marker), *NME1* (activation marker), *PRF1* (cytotoxic marker), and *IFNG*. Expression values are normalized for quantitative comparison within each dataset (i.e., column), but not across datasets.

A hallmark of tumor-associated T cells is a state of hyporesponsiveness or functional exhaustion, marked by persistent expression of surface inhibitory markers including PD-1, CTLA4, LAG3, TIM3 and others, many of which are expressed following T cell activation^17,50,51^. Some of these molecules (PD-1, CTLA4) are important targets for immunotherapy to promote anti-tumor immunity^52–56^. We compared expression of exhaustion and functional markers across healthy and tumor-associated T cells (Extended Data Fig. 6; Supplementary Fig. 7, 8). Tumor-associated CD8^+^T cells expressing exhaustion markers across all four tumor types project onto activated CD8^+^T cells in our map, and express genes within the Cytokine module (*CCL3*, *CCL4*, *XCL1*, *XCL2*, and *IFNG*, Supplementary Fig. 8), and to a lesser extent Cytotoxic module (Extended Data Fig. 6). Interestingly, a subset of these tumor-associated CD8^+^T cells, but not healthy T cells, express high levels of *MKI67*, associated with proliferating cells and other cell cycle control markers (Extended Data Fig. 6, Supplementary Fig. 8). Therefore, tumor-associated T cells expressing exhaustion markers also express genes associated with normal CD8 effector T cell function and ongoing proliferation.

## DISCUSSION

Human T cells persist in distinct anatomic sites, maintain protective immunity and surveillance, and are key targets for immune modulation in tumor immunotherapy, transplantation, and autoimmunity. Here, we used scRNA-seq profiling of resting and TCR-stimulated T cells from blood, lymphoid and mucosal tissues to generate a reference map of human T cells and understand how T cell homeostasis and function are related to the tissue site. Our findings demonstrate fundamental differences between T cells from tissues and blood, but similar functional and activation states across sites that are intrinsic to lineage; human CD4 T cell activation is defined by response to cytokines and proliferation while CD8 T cells are defined by effector function. We further demonstrate that this high resolution map of T cell homeostasis and activation across sites, lineages, and individuals can serve as a new baseline for defining human T cell states in disease.

The study of healthy human T cells has largely focused on blood, while the majority of T cells persist in diverse lymphoid, mucosal and barrier sites^8,57^. Human tissue T cells are largely memory subsets, comprising tissue-resident (TRM) and non-resident (TEM, TCM) populations; TRM predominate in mucosal sites, while TEM are found in spleen, LN and BM^13,33,58^. The relationship of these tissue-localized TEM to blood TEM has been unclear. Profiling using scRNAseq enabled unambiguous assessment of T cell-intrinsic differences in tissue versus blood T cells. We show that TEM from all sites examined (LG, LN, BM) exhibit fundamental changes in expression of cytoskeletal, cell-matrix interaction, and proliferative genes, indicating alterations in cellular structure. These tissue-intrinsic expression patterns are distinct from previously identified TRM-associated genes^13^, and are lowly expressed in T cells from blood. Whether T cells require these markers to localize within the tissue architecture, and/or their loss of expression enables egress to circulation remains to be established.

Our results reveal conserved functional states for human blood and tissue-derived T cells. CD8^+^T cells segregate into two major effector subsets based on expression of genes involved in cellular cytotoxicity (Cytotoxic module) and myriad cytokines and chemokines (Cytokine module). These predominant effector states within activated human CD8^+^T cells are consistent with results showing that mouse CD8^+^T cell activation triggers an effector differentiation program^59,60^. We identified two major activation states that were not associated with effector function: one associated with proliferation and IL-2 production, and a second, CD4-enriched state characterized by induction of a panoply of IFN-responsive genes including IFIT3, MX1, IRF7, and others. Induction of this IFN-response state is due to TCR-mediated IFN-γ production (likely autocrine responses), and appears as a kinetic intermediate early after CD4^+^T cell activation that subsequently shuts down upon induction of the proliferative program. We propose that the IFN-responsive state for human CD4^+^T cells may serve an autoregulatory function to temper high IFN levels produced by predominant memory responses, and ongoing responses to persistent viruses.

This scRNA-seq analysis provides a high resolution map for human T cells from which to define T cell states in disease. We demonstrate this approach by projecting T cell profiles from human tumors onto our reference map. We identify predominant CD8^+^ effector populations, Tregs, and resting (but not activated) CD4^+^T cells in datasets derived from diverse tumor types (breast, lung, skin, colon). Interestingly, the tumor-associated CD8^+^T cells exhibited transcriptional features similar to healthy activated CD8^+^T cells including expression of multiple effector molecules such as perforin, IFN-γ and chemokines. We also examined the expression of multiple markers associated with exhaustion, a functionally hyporesponsive state found in tumor-infiltrating T cells targeted by checkpoint blockade immunotherapies^53,55,61^. Interestingly, exhaustion markers were upregulated in activated T cells in both healthy and tumor tissues expressing CD8-associated cytokines. Moreover, subsets of these CD8^+^T cells in all four tumors expressed higher levels of proliferation markers compared to healthy T cells, consistent with a recent report that T cells expressing exhaustion markers in melanoma exhibit aberrant proliferation^62^. This analysis can therefore enable precise identification of features of resting and activated T cells that are associated with tissues, activation and disease.

In summary, our high resolution analysis of human T cells across sites, lineages, and activation states provides insights into human T cell adaptations to tissues and their intrinsic activation properties. This novel reference map can serve as a valuable resource for the study of human T cell immunity in disease, immunotherapies, vaccines and infections, and for diagnosing, screening and monitoring immune responses.

## ONLINE METHODS

### Acquisition of Human Tissues and Blood

We obtained human tissues from deceased, brain-dead donors at the time of organ acquisition for clinical transplantation through an approved research protocol and MTA with LiveOnNY, the organ procurement organization for the New York metropolitan area. Obtaining tissue samples from deceased organ donors does not qualify as “human subjects” research, as confirmed by the Columbia University Institutional Review Board (IRB). Donors were free of chronic disease, cancer and chronic infections such as Hepatitis B, C and HIV. Clinical and demographic data regarding organ donors used in this study are summarized in Supplementary Table 1. We obtained peripheral blood from healthy consenting adult volunteers by venipuncture, through an protocol approved by the Columbia University IRB.

### Isolation and Stimulation of T cells for Single-Cell RNA-seq

Tissues acquired from donors were maintained in cold saline during transport to the laboratory, typically within 2-4 hours of procurement. We isolated mononuclear cells from donor lungs, lung lymph nodes and bone marrow as previously described ^11,63^. Briefly, we flushed the left lobe of the lungs with cold complete medium (RPMI 1640, 10% FBS, 100 U/ml penicillin, 100 μg/ml streptomycin, 2 mM L-glutamine) and isolated lymph nodes from the hilum, near the intersections of major bronchi and pulmonary veins and arteries, removing all fat. To obtain mononuclear cell suspensions, hilar lymph nodes and the left lateral basal segment of the lung were mechanically processed using a gentleMACS tissue dissociator (Miltenyi Biotec), enzymatically digested (complete medium with 1 mg/ml collagenase D, 1 mg/ml trypsin inhibitor and 0.1 mg/ml DNase for 1 hour at 37°C in a mechanical shaker) and centrifuged on a density gradient using 30% Percoll Plus (GE Healthcare). We aspirated bone marrow from near the anterior superior iliac crest. For bone marrow and peripheral blood, we isolated mononuclear cells by density gradient centrifugation using Lymphocyte Separation Medium (Corning). Next, we enriched single cell suspensions from all tissues and blood for untouched CD3+ T cells using magnetic negative selection (MojoSort Human CD3+ T cell Isolation Kit; BioLegend). To eliminate any dead cells prior to stimulation, we used a dead cell removal kit (Miltenyi Biotec). We cultured 0.5 – 1 × 10^6^ CD3^+^ enriched cells from each donor tissue for 16 hours at 37°C in complete medium, with or without TCR stimulation using Human CD3/CD28 T Cell Activator (STEMCELL Technologies). After stimulation, we removed dead cells as above before preparing cells for single-cell RNA-seq.

### Single-Cell RNA-seq

We loaded single-cell suspensions into a Chromium Single Cell Chip (10x Genomics) according to the manufacturer’s instructions for co-encapsulation with barcoded Gel Beads at a target capture rate of ~5,000 individual cells per sample. We barcoded the captured mRNA during cDNA synthesis and converted the barcoded cDNA into pooled single-cell RNA-seq libraries for Illumina sequencing using the Chromium Single Cell 3’ Solution (10x Genomics) according to the manufacturer’s instructions. We processed all of the samples for a given donor simultaneously with the Chromium Controller (10x Genomics) and prepared the resulting libraries in parallel in a single batch. We pooled all of the libraries for a given donor, each of which was barcoded with a unique Illumina sample index, for sequencing in a single Illumina flow cell. All of the libraries were sequenced with an 8-base index read, a 26-base read 1 containing cell-identifying barcodes and unique molecular identifiers (UMIs), and a 98-base read 2 containing transcript sequences on an Illumina HiSeq 4000. Cell counts and transcript detection rates are summarized in Supplementary Table 2.

### Single-Cell RNA-seq Data Processing

Prior to gene expression analysis, we corrected the raw sequencing data for index swapping, a phenomenon that occurs during solid-phase clonal amplification on the Illumina HiSeq 4000 platform and results in cross-talk between sample index sequences. We corrected index swapping using the algorithm proposed by Griffiths *et al* ^64^. First, we aligned the reads associated with each sample index to GRCh38 (GENCODE v.24) using STAR v.2.5.0 after trimming read 2 to remove 3’ poly(A) tails (> 7 A’s) and discarding fragments with fewer than 24 remaining nucleotides as described in Yuan *et al* ^65^. For each read with a unique, strand-specific alignment to exonic sequence, we constructed an address comprised of the cell-identifying barcode, unique molecular identifier (UMI) barcode, and gene identifier. Next, we counted the number of reads associated with each address in each sample. Because of index swapping, we found that some addresses occurred in multiple samples at much higher frequencies than one would expect by chance. For the vast majority of addresses, there was a single sample containing most of the associated reads. If >80% of reads for a given address were associated with a single sample (e.g. a single index sequence), we kept all of the reads corresponding to that address in that sample and removed all of the reads associated with that address from all other samples^64^. We also identified addresses for which no sample contained >80% of the corresponding reads and removed all of these reads from all samples. After correcting for index swapping, we collapsed amplification duplicates using the UMIs and corrected errors in both the cell-identifying and UMI barcodes to generate a preliminary matrix of molecular counts for each cell as described previously ^65^.

We filtered the cell-identifying barcodes to avoid dead cells and other artifacts as described in Yuan *et al* ^65^. Briefly, we removed all cell-identifying barcodes where >10% of molecules aligned to genes expressed from the mitochondrial genome or for which the ratio of molecules aligning to whole gene bodies (including introns) to molecules aligning exclusively to exons was >1.5. Finally, we also removed cell-identifying barcodes for which the average number of reads per molecule or average number of molecules per gene deviated by >2.5 standard deviations from the mean for a given sample.

### Computational Identification of T Cells

As described above, we used negative selection to experimentally remove as many non-T cells as possible from our single-cell suspensions. This procedure was imperfect, and so all of our samples inevitably contained some non-T cells (average T cell purity was ~80%). Thoroughly removing non-T cells from the data set is complicated by technical issues such as molecular cross-talk, multiplet capture, and a broad coverage distribution. We developed a procedure to remove non-T cells that accounts for these issues by identifying both individual cells and clusters of cells that are enriched in expression of a blacklisted gene set that is highly specific to contaminating cell types.

We began by clustering the single-cell profiles within each sample using a pipeline that we reported previously ^65,66^. Briefly, we identified highly variable genes that are likely markers of specific subpopulations by normalizing the molecular counts for each cell to sum to one, ordering all genes by their normalized expression values, and computing a drop-out score *ds*_*g*_ for each gene *g* defined as:

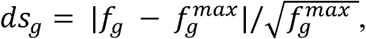

where *f_g_* is the fraction of cells in which we detected *g* and 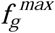 is the maximum *f*_*g*_ in a 25-gene rolling window centered on *g*. We selected genes with *ds*_*g*_ > 0.15 or with *ds*_*g*_ > 6σ_*ds*_ + < *ds*_*g*_ >, where σ_*ds*_ and < *ds*_*g*_ > are the standard deviation and mean of the dropout score distribution. Using these genes, we computed a cell-by-cell Spearman’s correlation, from which we constructed a k-nearest neighbor’s graph (k=20) and used this as input for the Phenograph^30^ implementation of Louvain clustering to identify cellular subpopulations.

Next, we used the pooled normalization approach described by Lun *et al* as implemented in the *scran* package with the *computeSumFactors* function to compute size factors for each cell ^67,68^. We supplied the *computeSumFactors* function with the cluster identifiers obtained from Phenograph to account for cell type-specific coverage differences. Using the resulting normalized expression profiles, we identified Phenograph clusters with positive enrichment of average *CD3D* and *TRAC* expression and labeled these clusters as T cell clusters (Supplementary Fig. S1). Within each sample, we conducted differential expression analysis between all pairs of T cell and non-T cell clusters via the Wilcoxon rank-sum test using the SciPy function *ranksums* and Benjamini-Hochberg corrected p-values with the StatsModels function *multipletests* in Python. Finally, we established an initial blacklist of genes that are highly specific to the non-T cell clusters by taking any gene with p < 0.001 and greater than 10 fold-enrichment in a non-T cell cluster for any of the above pairwise comparisons in any sample. To refine the blacklist and avoid including genes that are specific to T cell subsets found in only a limited set of samples or clusters, we also generated a whitelist of genes with positive enrichment in any T cell cluster. We removed any member of this whitelist from the initial blacklist to produce a final, refined blacklist containing 744 genes highly specific to contaminating cell types (Supplementary Table 3). As expected, genes on the final blacklist included markers of epithelial cells, dendritic cells, mast cells, B cells, neutrophils, and red blood cells.

We used the blacklist to remove cells from the T cell clusters that are either improperly clustered (unlikely to be T cells) or potentially multiplets (a cell-identifying barcode co-encapsulated both T cells and non-T cells). Importantly, because of molecular cross-talk in scRNA-seq libraries from PCR recombination, we only considered a cell to be expressing a blacklisted gene if the average number of reads supporting the detected molecules was above a certain threshold. This threshold depends on the average depth to which we sequenced the libraries in a given sample. The distributions of the number of reads-per-molecule are generally bimodal for a given sample. We assume that the mode with lower read counts per molecule arises from PCR recombination in which a molecule originating from one cell receives the cell-identifying barcode of a different cell at an intermediate point in PCR, thereby resulting in a detected molecule supported by an unusually small number of reads (i.e. amplicons). We therefore considered the sample *h* with the highest coverage (and therefore the clearest separation between the two modes) and took the minimum point between the two modes in the reads-per-molecule distribution to be the threshold number of reads per molecule, *T*_*h*_, below which a detected molecule would be considered to arise from cross-talk. We extrapolated a reads-per-molecule threshold for each of the other samples 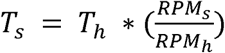, where *RPM*_*s*_ is the average number of reads per molecule detected in sample *s*.

Finally, for each cell *c* in a sample with threshold *T*_*s*_, we computed *b*_*c*_, the per-cell fraction of blacklisted genes detected with an average number of reads per molecule above *T*_*s*_. As expected, *b*_*c*_ was typically bimodally distributed within each sample (Supplementary Fig. 1e). The vast majority of cells in the lower mode were in the T cell clusters described above, while the high mode was composed mainly, but not exclusively, of cells from non-T cell clusters (Supplementary Fig. 1e). In each sample, we fit a Gaussian to *b*_*c*_’s distribution across cells assigned to T cell clusters and established a threshold at two standard deviations above the fitted mean. We considered any cell with *b*_*c*_ above this threshold and any cell that clustered among the non-T cell clusters to be a non-T cell and discarded these cells from all downstream analysis.

### Course-Grained Clustering of Merged T Cells from Each Donor

Once we had identified the T cells from each sample using the methodology described above, we merged resting and activated samples from all of the tissues in each donor and clustered the T cells from the two donors separately to generate Fig. 1b,c. We used the methodology described above to identify a set of highly variable genes for each sample (including the blood samples), and then merged those sets to generate a large list of 315 highly-variable genes (Supplementary Table 4) with which we clustered the merged samples from both donors. We computed Louvain clusters from the two merged data sets with k = 12 and a minimum cluster size of 100 cells using a k-nearest neighbors graph constructed from the Spearman’s correlation matrix calculated using the 315 highly variable genes. We used the Python implementation Uniform Manifold Approximation and Projection (UMAP)^31^ to produce the two-dimensional projections shown in Fig. 1b,c. To obtain *CD4*/*CD8* ratios for each cluster, we first computed the expression level of *CD4* and *CD8A* in each cell using the normalized counts from *computeSumFactors* as described above. For both *CD4* and *CD8A*, we then computed the average log_2_(normalized counts + 1) for each cluster and normalized this value by the average log_2_(normalized counts + 1) for all cells. We then took the log-ratio of these values for *CD4* and *CD8A* to generate Fig. 1b, where all the cells in each cluster are labeled with the cluster’s log-ratio.

### Blood Projection Analysis

To project the data obtained from blood T cells onto the tissue-derived profiles from each organ donor, we first merged the scRNA-seq profiles from both blood donors. We note that the scRNA-seq data from blood were subjected to the same computational procedure described above for eliminating non-T cell profiles. We used the same highly variable gene set (Supplementary Table 4) that was used in the original UMAP model of each organ donor to compute a Spearman’s correlation matrix between the blood and tissue profiles. We then projected the blood T cell profiles onto the UMAP models for each of the two organ donors using the *transform* function in UMAP. We note that the organ donor UMAP models used for this analysis are slightly different from what appears in Fig. 1b,c, because a small number of genes in the highly variable gene set were eliminated due to lack of expression in the blood. We also note that a small modification to the UMAP source code was needed to accommodate the use of Spearman’s correlation as a similarity metric.

To generate the cell number heatmaps in Fig. 2 and Supplementary Fig. 3, we first computed a centroid position in the UMAP embedding for each cluster-tissue combination in the tissue data based on the Louvain clustering described above for Fig. 1b,c. For example, for Donor 2, we computed the average position of lung-, bone marrow-, and lymph node-derived cells in the first cluster (CD4 Rest 1). We then identified the nearest cluster-tissue combination for each cell in the blood samples based on the Euclidean distance between a given blood-derived cell’s position in the UMAP model (following projection of the blood data onto the tissue UMAP model) and each cluster-tissue centroid position. The heatmaps summarize the results of these calculations, providing the number of blood-derived cells that are closest to each cluster-tissue combination in the organ donor data.

### Comparison of Tissue and Blood Effector Memory T Cells

To identify a tissue-specific T cell signatures, we compared the expression profiles of effector memory cells from resting LG, BM, and LN T cells from the two tissue donors to resting blood T cells from the two blood donors. We found *CCL5* to be an extremely highly expressed marker of effector memory cells that exhibited strong anti-correlation with *SELL*, a marker of non-effector memory cells, in all of our resting samples (Supplementary Fig. 4a). We also found that the average number of reads per molecule for *CCL5* was bimodally distributed, consistent with spurious detection of *CCL5* in a population of cells due to PCR recombination (Supplementary Fig. 4b). For each sample, we used the point between these two modes where the probability density was minimal as a threshold for the minimum average number of reads per molecule of *CCL5* required for a cell to be considered positive for *CCL5*. For each sample, we normalized the matrix of molecular counts for the *CCL5*^+^ effector memory T cells using the *computeSumFactors* function in *scran* to compute size factors for each cell^67,68^. For each tissue site, we then identified differentially expressed genes for all four pairwise comparisons of resting tissue to resting blood *CCL5*^+^ T cells (tissue donor 1 vs. blood donor A, tissue donor 2 vs. blood donor A, etc.) using the Wilcoxon rank-sum test with the SciPy function *ranksums* and computed Benjamini-Hochberg corrected p-values with the StatsModels function *multipletests* in Python after removing genes from the blacklist described above (Supplementary Table 3). For each tissue, we took all genes with p_adj_ < 0.05 and fold-change > 2 in all four pairwise comparisons to comprise a tissue-specific effector memory T cell signature (Fig. 3).

Next, all of the genes in the tissue-specific effector memory T cell signature and computed the average normalized expression of the resulting gene set to obtain Fig. 3d. Z-scored normalized expression for each of these genes appears in the heatmap in Fig. 3e for each site / donor, which also includes a set of blood T cells with outlier expression of the tissue-enriched gene signature (blood T cells with average expression within one standard deviation of that of the tissue T cells as indicated by the dashed line in Fig. 3d).

### Single-cell Hierarchical Poisson Factorization and Transcriptional Module Analysis

We applied Single-cell Hierarchical Poisson Factorization (scHPF), a method that we recently reported for *de novo* discovery of gene expression signatures in scRNA-seq data, to the merged activated and resting cells for each tissue and donor^66^. Given a molecular count matrix, scHPF identifies a small number of latent factors that explain both continuous and discrete expression patterns across cells. Each gene has a score for each factor, quantifying the gene’s contribution to the associated expression pattern. Likewise, each cell assigns a score to each factor, which reflects the contribution of the factor to the observed expression in the cell.

We applied scHPF to each tissue and blood sample after merging their respective resting and activated datasets. We considered only genes with GENCODE protein coding, T cell receptor constant or immunoglobulin constant biotypes, excluded genes on the previously described blacklist, and removed genes detected in fewer than 0.1% of cells in a given merged dataset. scHPF was run with default parameters for seven values of *K*, the number of factors, equal to all values between 6-12, inclusively. This resulted in seven candidate scHPF models per merged dataset. We then selected *K* (and a corresponding fitted model) to avoid factors with significant overlap in their gene signatures. For each dataset and value of *K*, we calculated *N*_*K*_: the maximum pairwise overlap of the 300 highest-scoring genes in each factor for the corresponding scHPF model. We considered overlap significant if p < 0.05 by a hypergeometric test with a population size equal to the number of unfiltered genes in the tissue sample and *N*_*K*_ observed successes. Finally, for each dataset, we selected the model with maximum *K* such that p>=0.05 (Supplementary Fig. 5). This resulted in eight factorizations: six from tissue donors (lung, bone marrow, and lymph node from each of two organ donors) and two factorizations from the blood of living donors. We defined each factors’ CD4/CD8 bias as the log2 ratio of its mean cell score in CD4^+^ and CD8^+^T cells.

To discover common patterns of expression across tissues and donors, we performed unsupervised clustering of all factors for tissue-derived cells. First, we calculated Pearson correlation on the union of the fifty highest and lowest scoring genes in each factor for each tissue factorization (2,291 genes total) using the Python pandas package’s *DataFrame.corr* function. Next, we hierarchically clustered the factor-factor correlation matrix using *scipy.cluster.hierarchy.linkage* with method=’average’ and *scipy.cluster.hierarchy.dendrogram* (Extended Data Fig. 4). This defined clusters of tightly correlated expression patterns, which we call expression modules. We focused on seven modules (out of nine) whose factors had mean pairwise correlations greater than 0.25. Most modules contained at least one factor from each tissue and donor. To identify the top genes in each module (Fig. 4a, Supplementary Table 7), we ranked genes by their mean gene score across all constituent factors. Finally, we noticed that the CD4 IFN response module contained two factors from Donor 2’s bone marrow; however, one of the two factors was far more tightly correlated with the rest of the factors in the module than the other. As the top genes in the module were nearly identical with and without the less tightly-correlated factor, we excluded it from the module in downstream analyses for clarity.

### Activation Trajectory Analysis

We used the factorizations described above to compute T cell activation trajectories by diffusion component analysis. We first converted the cell score matrix obtained from the factorization of each resting/activated merged tissue or blood sample into a cell-by-cell Euclidean distance matrix. We then extracted the distance submatrices corresponding to the CD4 and CD8 clusters in each sample as defined from the merged analysis of all samples from each donor described above. We used the two resulting distance submatrices to compute diffusion components for CD4 and CD8 activation with the C++ Accelerated Python Diffusion Maps Library (DMAPS) with a kernel bandwidth of four. The diffusion maps shown in Fig. 4b-e each show the first two diffusion components which we define as the two diffusion eigenvectors with the second-and third-highest eigenvalues scaled by the diffusion eigenvector with the largest eigenvalue.

### Flow Cytometry, Intracellular Staining and Proliferation Assays

To evaluate the expression of T cell surface markers by flow cytometry, we incubated tissue and blood cell suspensions with Human TruStain FcX (BioLegend) and stained with following fluorochrome-conjugated antibodies: CD3 (UCHT1, BD Biosciences; OKT3, BioLegend), CD4 (SK3, BD Biosciences; SK3, Tonbo Biosciences), CD8 (SK1, BioLegend; RPA-T8, BD Biosciences), CCR7 (G043H7; BioLegend), CD45RA (HI100; BioLegend), CD25 (BC96; BioLegend), CD127 (A019D5; BioLegend), CD69 (FN50; BioLegend), CD103 (Ber-ACT8; BioLegend), CD45 (HI30; BioLegend), and Fixable Viability Dye eFluor 780 (eBioscience). For stimulation/proliferation assays, we magnetically enriched for CD3+ T cells from single cell suspensions, stained cells with Cell Proliferation Dye eFluor 450 (eBioscience), and cultured cells for up to 120 hours with or without TCR stimulation as above. At indicated time points, we performed intercellular staining of NME1 (11615-H07E; Sino Biological) using a Foxp3/Transcription Factor Staining Buffer Kit (Tonbo Biosciences) for fixation and permeabilization of cells according to manufacturer’s instructions. We acquired cell fluorescence data using a BD LSR II flow cytometer and used FCS Express (De Novo Software) for analysis. The results are summarized in Supplementary Fig. 2 and the gating strategy is shown in Supplementary Fig. 9.

### Gene Expression Kinetics by Quantitative Real-Time PCR

We isolated mononuclear cells from peripheral blood, magnetically enriched for CD3+ T cells, and sorted live CD4+ and CD8+ T cells (gated for singlets, FSC^low^SSC^low^, CD45^+^ and Viability Dye^-^) using a BD Influx cell sorter. Sorted cells were cultured in complete medium with or without anti-CD3/anti-CD28 stimulation as above for up to 72 hours. For dissecting the contribution of type I and type II IFN signaling to gene expression, cells were pre-incubated with Human Type 1 IFN Neutralizing Antibody Mixture (PBL Assay Science, Cat# 39000-1) according to manufacturer’s instructions, or 1 ug/mL of both anti-IFNγ (R&D Systems, MAB285, clone # 25718) and anti-IFNγR1 (R&D Systems, MAB6731, clone # 92101). As a control, CD4+ T cells were activated with 1000 units/mL of recombinant human IFNα2 (PBL Assay Science, Cat#11101-1) or 10 ng/mL recombinant human IFNγ (Peprotech, Cat# 300-02). We harvested resting and activated CD4+ and CD8+ T cells at indicated time points and extracted RNA using a RNeasy Micro Kit (Qiagen) with on-column DNase digestion. We converted RNA to cDNA via SuperScript IV VILO Master Mix (Invitrogen) and performed quantitative real-time PCR (qPCR) on a Viia 7 Real-Time PCR system (Applied Biosystems) using TaqMan Gene Expression Assays (NME1 Hs00264824_m1; IL2RA Hs00907777_m1; IFIT3 Hs00155468_m1; TBP Hs00427620_m1) and TaqMan Fast Advanced Master Mix, all from ThermoFisher Scientific. qPCR reactions were set up according to manufacturer’s instructions and fold changes between stimulated and unstimulated cells at each time point were calculated using the ∆∆ cycle threshold method in ExpressionSuite Software (ThermoFisher Scientific) with TBP as a reference gene.

### Tumor-Associated T Cell Projection Analysis

We projected scRNA-seq profiles of tumor-associated T cells from four different tumor types onto a UMAP embedding of resting and activated T cells from across our combined tissue and blood data set using the methods described above for projecting blood T cells onto embeddings of the tissues. Briefly, we used the highly variable gene set from Supplementary Table 4 to generate a UMAP embedding of our tissue/blood data set from a Spearman’s correlation matrix. We then projected the tumor-associated T cell profiles onto this embedding using the *transform* function in UMAP. Tumor-associated T cells from non-small cell lung cancer (NSCLC)^49^ and breast cancer (BC)^48^, which were profiled using the 10x Genomics Chromium platform, were obtained from https://gbiomed.kuleuven.be/scRNAseq-NSCLC and GEO accession GSE114724 (samples BC09, BC10, and BC11), respectively. For these two data sets, we used the UMI-corrected molecular counts provided by the authors. T cells from colorectal cancer (CRC)^47^ and melanoma (MEL)^27^, which were profiled using SMART-seq, were obtained from GEO accessions GSE108989 and GSE120575 (pre-treated samples only). For these two data sets, we used the TPM values provided by the authors. We note that the tissue/blood embedding was re-computed for each projection and is therefore slightly different in each case because not all of the processed data sets from the tumor studies contained all of the genes in Supplementary Table 4.

The resulting projections are displayed in Fig. 6 in three different ways. In the top row, the projections are displayed as contour plots of estimated probability density (kernel density estimates) with a maximum of 14 contours. In the second row, we used a hexbin two-dimensional histogram of the number of cells in each bin with the colorbars normalized such that the intensity can be compared across samples (e.g. scaled so that the melanoma projection can be compared to the CRC projection). Finally, we also show where individual tumor-associated T cells project in subsequent rows along with gene expression values for several key markers. In Extended Data Fig. 4, we show the average expression of several canonical exhaustion markers in individual cells. The markers used for this analysis were *PDCD1*, *CTLA4*, *LAG3*, *LAYN*, *TIM-3*, *CD244*, and *CD160*.

### Data Availability

The scRNA-seq data are available on the Gene Expression omnibus (GEO) under accession number GSE126030.

### Code Availability

The computer code for scHPF is freely available at www.github.com/simslab/scHPF.

## Acknowledgements

This work was supported by the US National Institutes of Health (NIH) (grant nos. AI128949, AI106697 to D.L.F.; AI128949, AI106697 and Chan Zuckerberg Initiative Pilot Projects for the Human Cell Atlas to P. A. Sims.). These studies were performed in the Columbia Center for Translational Immunology (CCTI) Flow Cytometry Core funded in part through an S10 Shared Instrumentation Grant from the NIH (grant no. S10RR027050), with the excellent technical assistance of S.-H. Ho. We gratefully acknowledge the generosity of the organ donor families and the Dr. Amy Friedman and the LiveOnNY transplant coordinators and staff for making this study possible.

## Author contributions

P.A.Sz. designed, executed and analyzed experiments; H.M.L. developed the scHPF module analysis approach, H.M.L and P.A.Si. performed computational analysis; T.E.S. obtained tissues from donors; M.M., M.E.S., processed tissues and optimized protocols, E.C.B. constructed and sequenced the scRNA-seq libraries, J.Y. and Y.L.C. optimized scRNA-seq experiments, P.D., and P.T. provided technical assistance; P.A.Sz., H.M.L., D.L.F., and P.A.Si. analyzed data, wrote and edited the manuscript; D.L.F. and P.A.Si. designed and coordinated the study.

## Extended Data Figure Legends

**Extended Data Figure 4.**
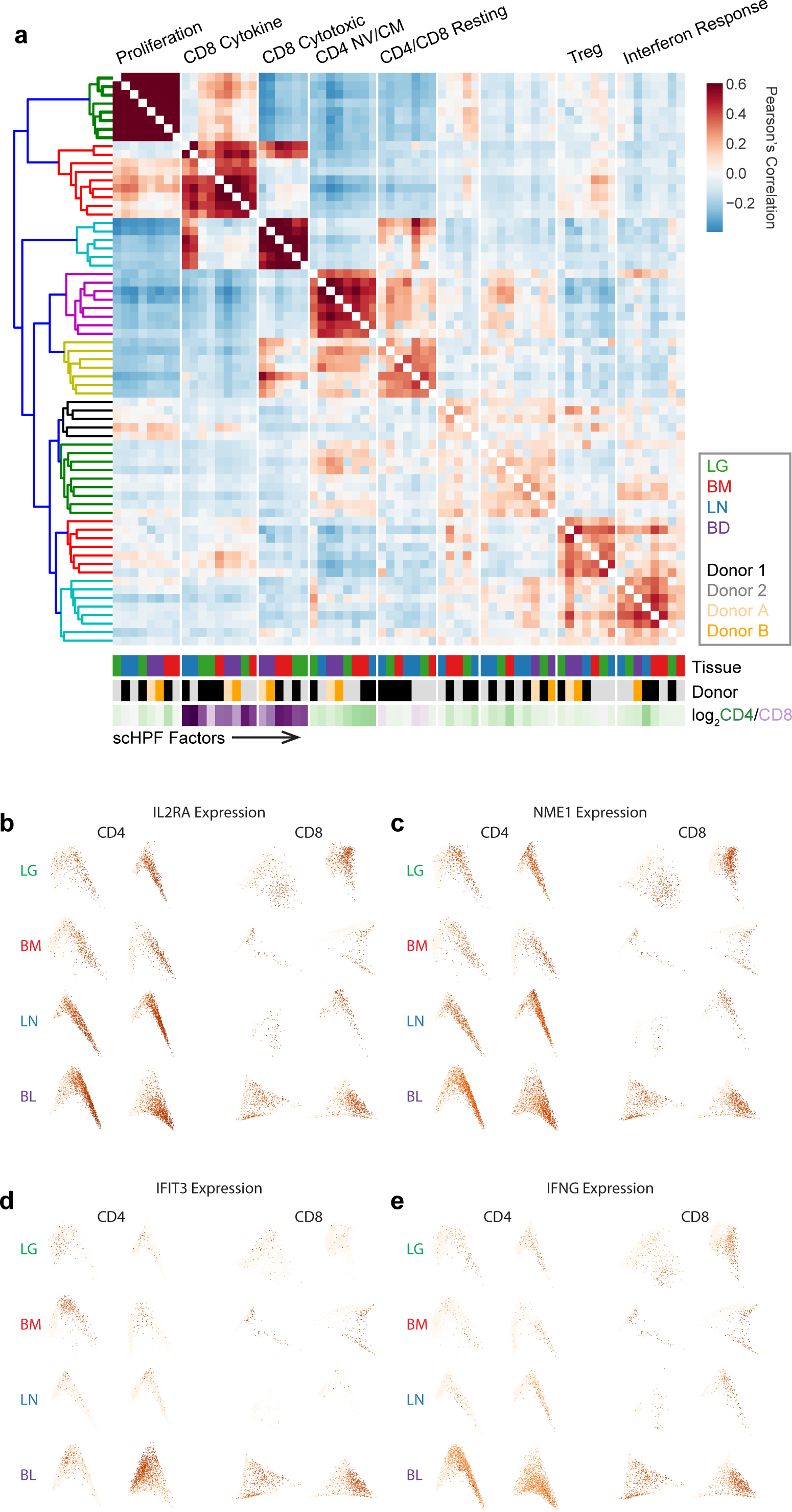
(a) scHPF was used to factorize scRNA-seq profiles of each tissue and blood sample independently after merging resting and activated T cells. Each matrix element in the heatmap is the pairwise Pearson’s correlation coefficient between the gene scores for a pair of factors computed across a set of high-and low-scoring genes (see Online Methods). The resulting modules were named based on the identities of their highest scoring genes (Supplementary Table 7) and the resting vs. activated status of the highest scoring cells. Lower color bars show the tissue and donor of origin, and the CD4/CD8 bias of cell scores for each factor. (b) Diffusion maps generated from scHPF factors (see Online Methods) for CD4^+^ and CD8^+^ T cells from each tissue and blood sample after merging resting and activated T cells with each cell colored by expression of *IL2RA*. (c) Same as (b) but colored by expression of *NME1*. (d) Same as (b) but colored by expression of *IFIT3*. (e) Same as (b) but colored by expression of *IFNG*.

**Extended Data Figure 6.**
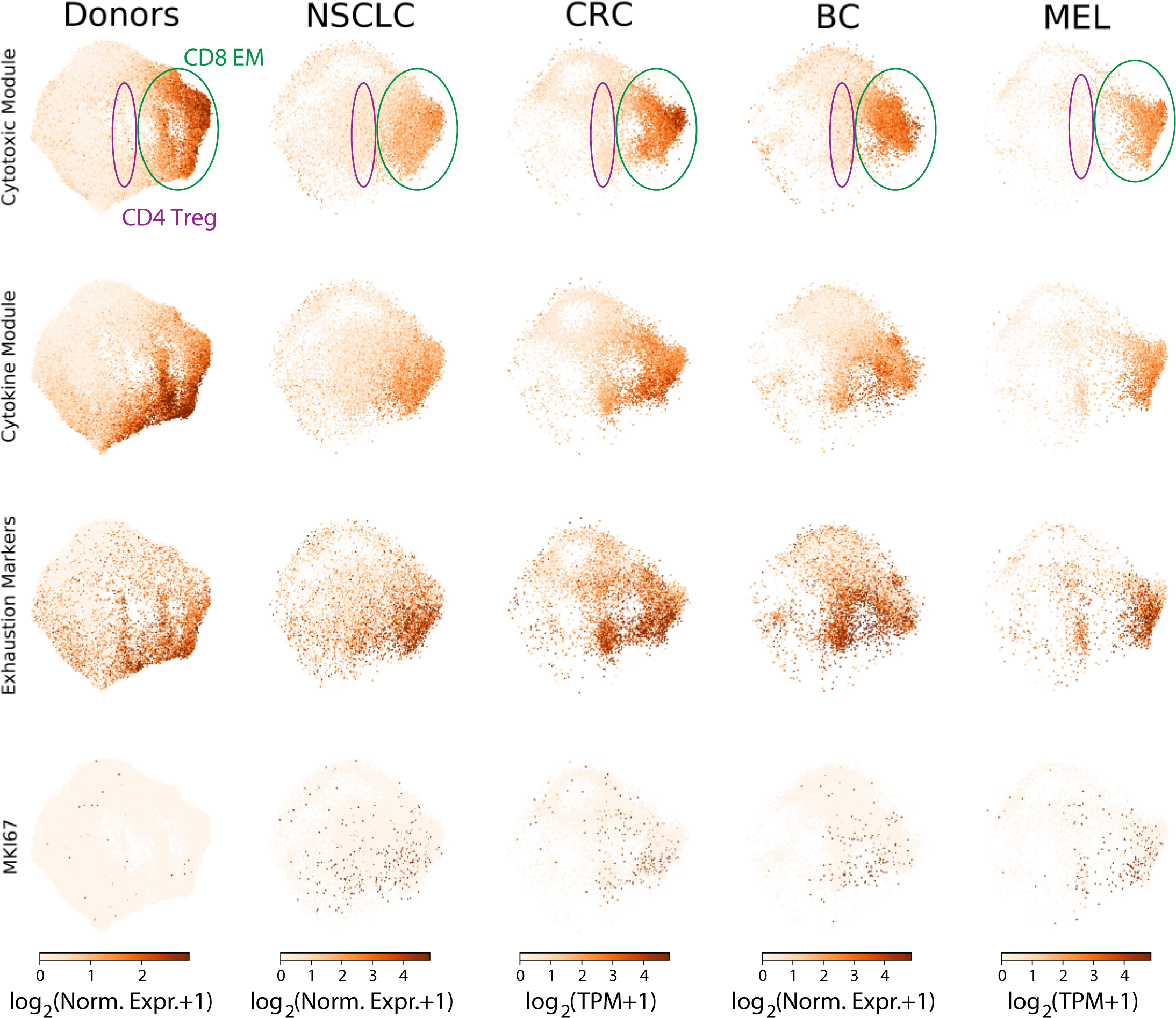
Individual cells in the UMAP embedding (far left column) for the entire healthy T cell dataset and UMAP projections (remaining four columns) for NSCLC, CRC, BC, and MEL tissue T cells colored by the average expression of the top 70 genes in the Cytotoxic module, the top 70 genes in the Cytokine module, the average expression of a set of exhaustion markers (*PDCD1*, *CTLA4*, *LAYN*, *LAG3*, *TIM-3*, *CD244*, and *CD160*), and expression of the proliferation marker *MKI67*. Note that these expression values are normalized so that they can be quantitatively compared within each dataset (within each column), but not across datasets.

## References

1. Sallusto, F., Lenig, D., Forster, R., Lipp, M. & Lanzavecchia, A. Two subsets of memory T lymphocytes with distinct homing potentials and effector functions. Nature 401, 708–712 (1999).

2. Teijaro, J.R., et al. Cutting edge: tissue-retentive lung memory CD4 T cells mediate optimal protection to respiratory virus infection. J Immunol 187, 5510–5514 (2011).

3. Mackay, L.K., et al. The developmental pathway for CD103(+)CD8+ tissue-resident memory T cells of skin. Nat Immunol 14, 1294–1301 (2013).

4. Schenkel, J.M. & Masopust, D. Tissue-Resident Memory T Cells. Immunity 41, 886–897 (2014).

5. Schenkel, J.M., et al. T cell memory. Resident memory CD8 T cells trigger protective innate and adaptive immune responses. Science 346, 98–101 (2014).

6. Park, S.L., et al. Local proliferation maintains a stable pool of tissue-resident memory T cells after antiviral recall responses. Nat Immunol 19, 183–191 (2018).

7. Wilk, M.M., et al. Lung CD4 Tissue-Resident Memory T Cells Mediate Adaptive Immunity Induced by Previous Infection of Mice with Bordetella pertussis. J Immunol (2017).

8. Ganusov, V.V. & De Boer, R.J. Do most lymphocytes in humans really reside in the gut? Trends Immunol 28, 514–518 (2007).

9. Carpenter, D.J., et al. Human immunology studies using organ donors: Impact of clinical variations on immune parameters in tissues and circulation. Am J Transplant 18, 74–88 (2018).

10. Thome, J.J.C., et al. Spatial Map of Human T Cell Compartmentalization and Maintenance over Decades of Life. Cell 159, 814–828 (2014).

11. Sathaliyawala, T., et al. Distribution and compartmentalization of human circulating and tissue-resident memory T cell subsets. Immunity 38, 187–197 (2013).

12. Miron, M., et al. Human Lymph Nodes Maintain TCF-1(hi) Memory T Cells with High Functional Potential and Clonal Diversity throughout Life. J Immunol 201, 2132–2140 (2018).

13. Kumar, B.V., et al. Human tissue-resident memory T cells are defined by core transcriptional and functional signatures in lymphoid and mucosal sites. Cell Reports 20, 2921–2934 (2017).

14. Nakayamada, S., Takahashi, H., Kanno, Y. & O’Shea, J.J. Helper T cell diversity and plasticity. Curr Opin Immunol 24, 297–302 (2012).

15. Kaech, S.M. & Wherry, E.J. Heterogeneity and cell-fate decisions in effector and memory CD8+ T cell differentiation during viral infection. Immunity 27, 393–405 (2007).

16. Wherry, E.J., et al. Molecular signature of CD8+ T cell exhaustion during chronic viral infection. Immunity 27, 670–684 (2007).

17. Wherry, E.J. T cell exhaustion. Nat Immunol 12, 492–499 (2011).

18. Zajac, A.J., et al. Viral immune evasion due to persistence of activated T cells without effector function. J Exp Med 188, 2205–2213 (1998).

19. Salgame, P., et al. Differing lymphokine profiles of functional subsets of human CD4 and CD8 T cell clones. Science 254, 279–282 (1991).

20. Yang, L., et al. IL-21 and TGF-beta are required for differentiation of human T(H)17 cells. Nature 454, 350–352 (2008).

21. Fromentin, R., et al. CD4+ T Cells Expressing PD-1, TIGIT and LAG-3 Contribute to HI V Persistence during ART. PLoS Pathog 12, e1005761 (2016).

22. Banga, R., et al. PD-1(+) and follicular helper T cells are responsible for persistent HIV-1 transcription in treated aviremic individuals. Nat Med 22, 754–761 (2016).

23. Papalexi, E. & Satija, R. Single-cell RNA sequencing to explore immune cell heterogeneity. Nat Rev Immunol 18, 35–45 (2018).

24. Griffiths, J.A., Scialdone, A. & Marioni, J.C. Using single-cell genomics to understand developmental processes and cell fate decisions. Mol Syst Biol 14, e8046 (2018).

25. De Simone, M., et al. Transcriptional Landscape of Human Tissue Lymphocytes Unveils Uniqueness of Tumor-Infiltrating T Regulatory Cells. Immunity 45, 1135–1147 (2016).

26. Stubbington, M.J.T., Rozenblatt-Rosen, O., Regev, A. & Teichmann, S.A. Single-cell transcriptomics to explore the immune system in health and disease. Science 358, 58–63 (2017).

27. Sade-Feldman, M., et al. Defining T Cell States Associated with Response to Checkpoint Immunotherapy in Melanoma. Cell 175, 998–1013 e1020 (2018).

28. Thome, J.J., et al. Longterm maintenance of human naive T cells through in situ homeostasis in lymphoid tissue sites. Sci Immunol 1, aah6506 (2016).

29. Granot, T., et al. Dendritic Cells Display Subset and Tissue-Specific Maturation Dynamics over Human Life. Immunity 46, 504–515 (2017).

30. Levine, J.H., et al. Data-Driven Phenotypic Dissection of AML Reveals Progenitor-like Cells that Correlate with Prognosis. Cell 162, 184–197 (2015).

31. McInnes, L. & Healy, J. UMAP: Uniform Manifold Approximation and Projection for Dimension Reduction. arXiv, arXiv:1802.03426 (2018).

32. Thome, J.J., et al. Early-life compartmentalization of human T cell differentiation and regulatory function in mucosal and lymphoid tissues. Nat Med 22, 72–77 (2016).

33. Hombrink, P., et al. Programs for the persistence, vigilance and control of human CD8+ lung-resident memory T cells. Nat Immunol 17, 1467–1478 (2016).

34. Swanson, B.J., Murakami, M., Mitchell, T.C., Kappler, J. & Marrack, P. RANTES production by memory phenotype T cells is controlled by a posttranscriptional, TCR-dependent process. Immunity 17, 605–615 (2002).

35. Levitin, H.M., et al. De novo Gene Signature Identification from Single-Cell RNA-Seq with Hierarchical Poisson Factorization. bioRxiv, 367003 (2018).

36. Zemmour, D., et al. Single-cell gene expression reveals a landscape of regulatory T cell phenotypes shaped by the TCR. Nat Immunol 19, 291–301 (2018).

37. Kondrack, R.M., et al. Interleukin 7 regulates the survival and generation of memory CD4 cells. J Exp Med 198, 1797–1806 (2003).

38. Tan, J.T., et al. IL-7 is critical for homeostatic proliferation and survival of naive T cells. Proc Natl Acad Sci U S A 98, 8732–8737 (2001).

39. Moon, C., et al. Aquaporin expression in human lymphocytes and dendritic cells. Am J Hematol 75, 128–133 (2004).

40. Boissan, M. & Lacombe, M.L. Learning about the functions of NME/NM23: lessons from knockout mice to silencing strategies. Naunyn Schmiedebergs Arch Pharmacol 384, 421–431 (2011).

41. Schoggins, J.W. & Rice, C.M. Interferon-stimulated genes and their antiviral effector functions. Curr Opin Virol 1, 519–525 (2011).

42. Schneider, W.M., Chevillotte, M.D. & Rice, C.M. Interferon-stimulated genes: a complex web of host defenses. Annu Rev Immunol 32, 513–545 (2014).

43. Dominguez, C.X., et al. The transcription factors ZEB2 and T-bet cooperate to program cytotoxic T cell terminal differentiation in response to LCMV viral infection. J Exp Med 212, 2041–2056 (2015).

44. Mackay, L.K., et al. Hobit and Blimp1 instruct a universal transcriptional program of tissue residency in lymphocytes. Science 352, 459–463 (2016).

45. Pearce, E.L., et al. Control of effector CD8+ T cell function by the transcription factor Eomesodermin. Science 302, 1041–1043 (2003).

46. Mariotto, A., Pavlova, O., Park, H.S., Huber, M. & Hohl, D. HOPX: The Unusual Homeodomain-Containing Protein. J Invest Dermatol 136, 905–911 (2016).

47. Zhang, L., et al. Lineage tracking reveals dynamic relationships of T cells in colorectal cancer. Nature 564, 268–272 (2018).

48. Azizi, E., et al. Single-Cell Map of Diverse Immune Phenotypes in the Breast Tumor Microenvironment. Cell 174, 1293–1308 e1236 (2018).

49. Lambrechts, D., et al. Phenotype molding of stromal cells in the lung tumor microenvironment. Nat Med 24, 1277–1289 (2018).

50. Barber, D.L., et al. Restoring function in exhausted CD8 T cells during chronic viral infection. Nature 439, 682–687 (2006).

51. Blackburn, S.D., et al. Coregulation of CD8+ T cell exhaustion by multiple inhibitory receptors during chronic viral infection. Nat Immunol 10, 29–37 (2009).

52. Yuan, J., et al. CTLA-4 blockade enhances polyfunctional NY-ESO-1 specific T cell responses in metastatic melanoma patients with clinical benefit. Proc Natl Acad Sci U S A 105, 20410–20415 (2008).

53. Sharma, P. & Allison, J.P. The future of immune checkpoint therapy. Science 348, 56–61 (2015).

54. Phan, G.Q., et al. Cancer regression and autoimmunity induced by cytotoxic T lymphocyte-associated antigen 4 blockade in patients with metastatic melanoma. Proc Natl Acad Sci U S A 100, 8372–8377 (2003).

55. Chen, R., et al. Anti-Programmed Cell Death (PD)-1 Immunotherapy for Malignant Tumor: A Systematic Review and Meta-Analysis. Transl Oncol 9, 32–40 (2016).

56. Robert, C., et al. Nivolumab in previously untreated melanoma without BRAF mutation. N Engl J Med 372, 320–330 (2015).

57. Farber, D.L., Yudanin, N.A. & Restifo, N.P. Human memory T cells: generation, compartmentalization and homeostasis. Nat Rev Immunol 14, 24–35 (2014).

58. Kumar, B.V., Connors, T.J. & Farber, D.L. Human T Cell Development, Localization, and Function throughout Life. Immunity 48, 202–213 (2018).

59. Kaech, S.M., Hemby, S., Kersh, E. & Ahmed, R. Molecular and functional profiling of memory CD8 T cell differentiation. Cell 111, 837–851 (2002).

60. Zhang, N. & Bevan, M.J. CD8(+) T cells: foot soldiers of the immune system. Immunity 35, 161–168 (2011).

61. McLane, L.M., Abdel-Hakeem, M.S. & Wherry, E.J. CD8 T Cell Exhaustion During Chronic Viral Infection and Cancer. Annu Rev Immunol (2019).

62. Li, H., et al. Dysfunctional CD8 T Cells Form a Proliferative, Dynamically Regulated Compartment within Human Melanoma. Cell (2018).

63. Thome, J.J., et al. Spatial map of human T cell compartmentalization and maintenance over decades of life. Cell 159, 814–828 (2014).

64. Griffiths, J.A., Richard, A.C., Bach, K., Lun, A.T.L. & Marioni, J.C. Detection and removal of barcode swapping in single-cell RNA-seq data. Nat Commun 9, 2667 (2018).

65. Yuan, J., et al. Single-cell transcriptome analysis of lineage diversity in high-grade glioma. Genome Med 10, 57 (2018).

66. Levitin, H.M., et al. De novo Gene Signature Identification from Single-Cell RNA-Seq with Hierarchical Poisson Factorization. bioRxiv (2018).

67. Lun, A.T., Bach, K. & Marioni, J.C. Pooling across cells to normalize single-cell RNA sequencing data with many zero counts. Genome Biol 17, 75 (2016).

68. Lun, A.T., McCarthy, D.J. & Marioni, J.C. A step-by-step workflow for low-level analysis of single-cell RNA-seq data with Bioconductor. F1000Res 5, 2122 (2016).

