## Supplementary Figures and Tables for "A single-cell reference map for human blood and tissue T cell activation reveals functional states in health and disease"

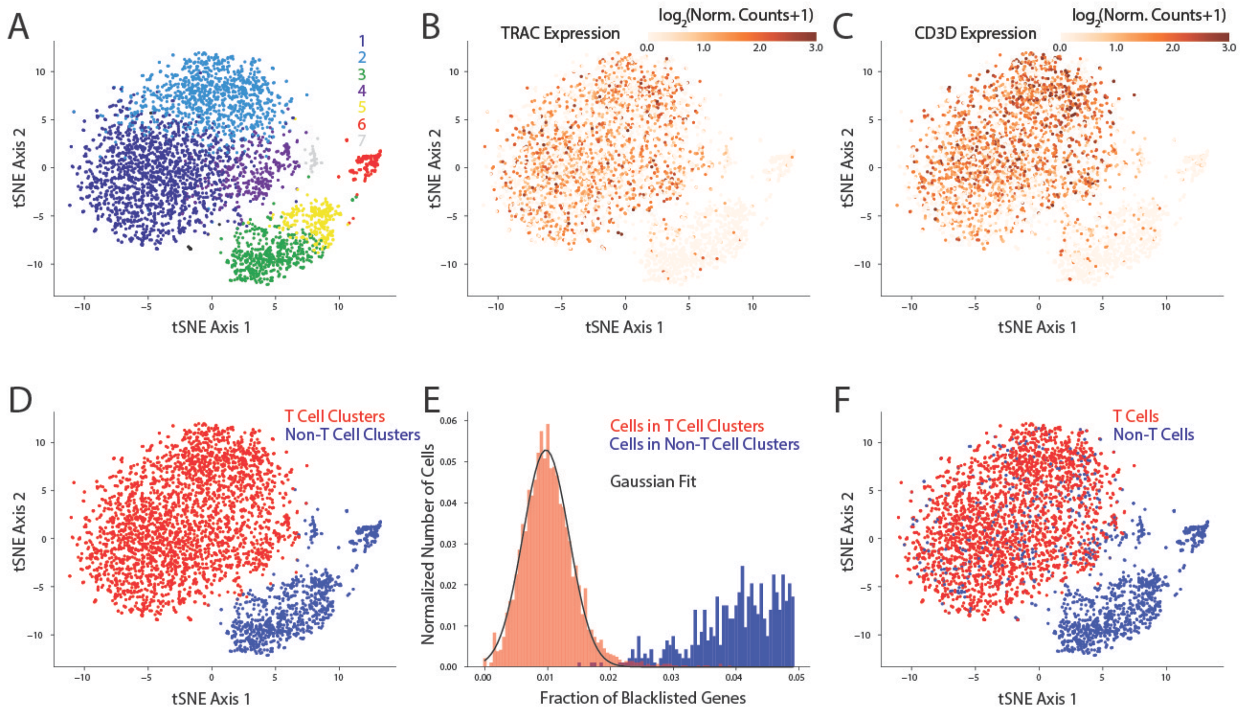


**Supplementary Figure 1. Example of computational methods for identifying T cells from single-cell RNA-seq data**. (A) tSNE projection scRNA-seq profiles of the resting cell sample from LG of Tissue Donor 1. Individual cells are colored by cluster membership identified using Phenograph (see Supplementary Methods). (B) Same as (A) with cells colored by expression of TRAC, a highly expressed marker of T cells. (C) Same as (A) with cells colored by expression of *CD3D*, a highly expressed marker of T cells. (D) Same as (A) with cells colored by whether they are members of clusters that are enriched in *CD3D* expression (red) or not (blue). (E) Histograms of the fraction of blacklisted genes detected per cell for cells in clusters that are enriched in *CD3D* expression (red) and cells that are not (blue). The blacklisted genes are identified by differential expression analysis between the red and blue cells across the entire data set (see Supplementary Methods). The black line is a Gaussian fit to the red histogram. (F) Same as (A) with cells colored by whether they are identified as T cells (red) or not (blue). All cells in clusters that are not enriched in *CD3D* are considered non-T cells. Cells in the clusters that are enriched in *CD3D* are considered non-T cells if the fraction of blacklisted genes detected exceeds two standard deviations above the mean of the Gaussian fit in E).

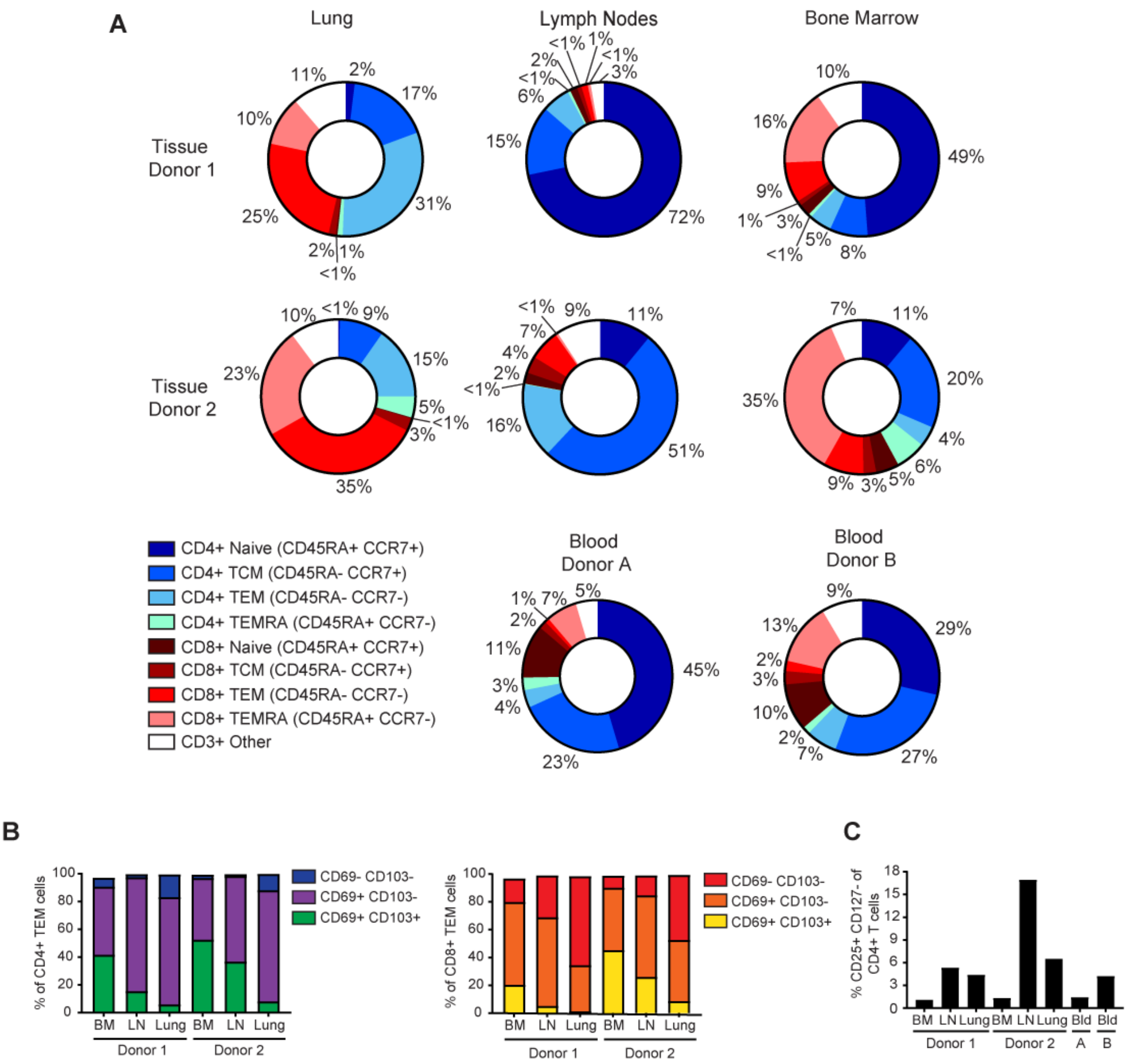


**Supplementary Figure 2. T cell subset phenotypes for tissue and blood donors.** (A) T cell subset composition of tissue and blood donors, showing frequency of Naïve (CD45RA+ CCR7+), central-memory (TCM; CD45RA- CCR7+) effector-memory (TEM; CD45RA- CCR7-), and terminal effector (TEMRA; CD45RA+ CCR7-) subsets for $CD4^+$ and $CD8^+$T cells. (B) Fraction of TEM cells expressing tissue resident memory (TRM) markers CD69 +/- CD103 by $CD4^+$ and $CD8^+$TEM cells from the two tissue donors. (C) Fraction of $CD4^+$T cells in each sample (both tissue and blood donors) that express a regulatory T cell (Treg) phenotype ($CD25^+$/$CD127^-$).

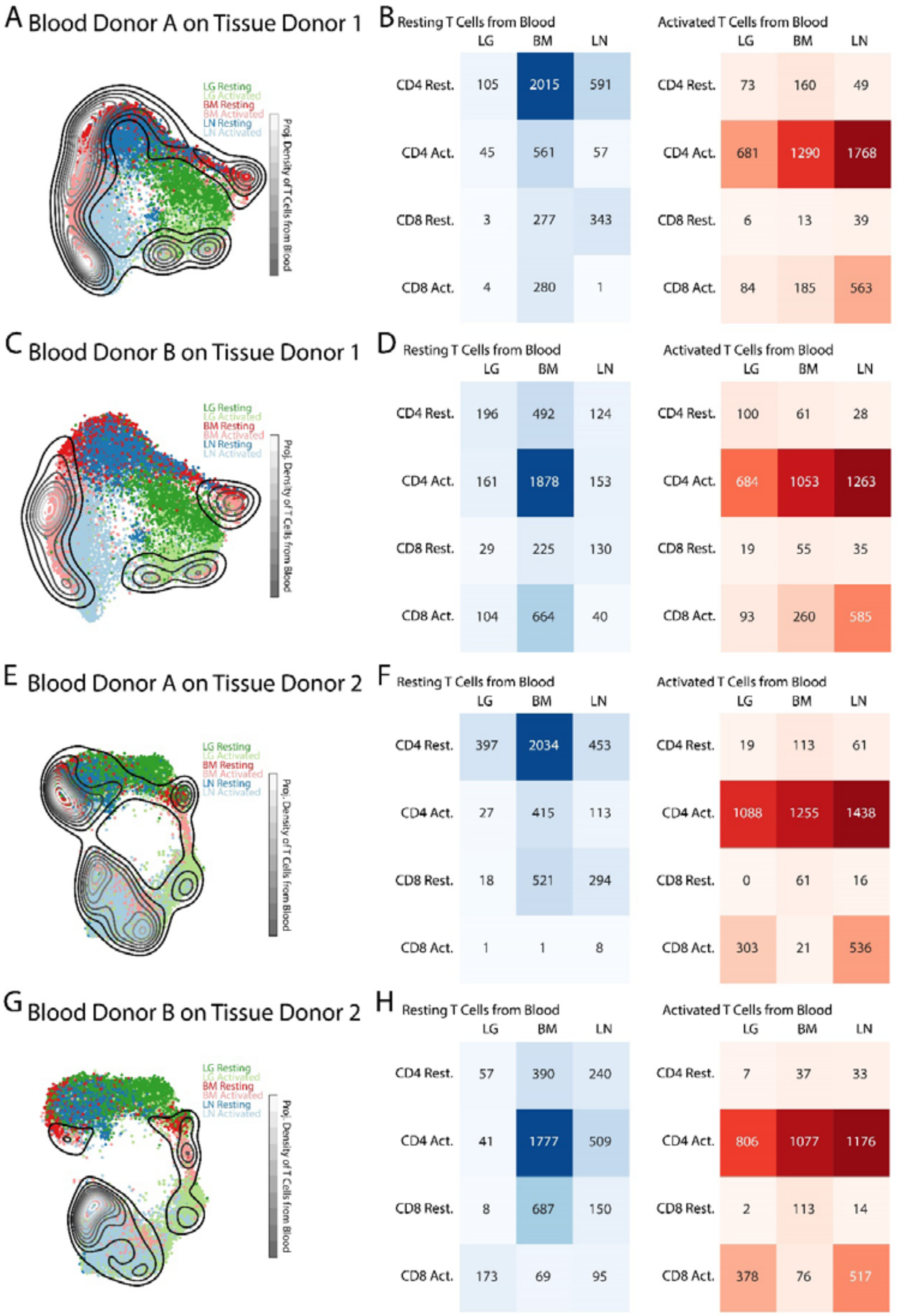


**Supplementary Figure 3. Projections of individual blood T cells onto UMAP embeddings of individual tissue T cells.** (A) UMAP embedding of T cells from tissue donor 1 colored by tissue, overlaid by a contour plot corresponding to the projection of merged resting and activated T cells from blood donor A onto the tissue embedding. (B) Heatmaps showing the number of blood donor A T cells that project most closely to each tissue-activation combination in the tissue donor 1 UMAP embedding. (C) Same as (A) for blood donor B. (D) Same as (B) for blood donor B. (E)

Same as (A) for tissue donor 2 and blood donor A. (F) Heatmaps showing the number of blood donor A T cells that project most closely to each tissue-activation combination in the tissue donor 2 UMAP embedding. G) Same as E) for blood donor B. H) Same as F) for blood donor B.

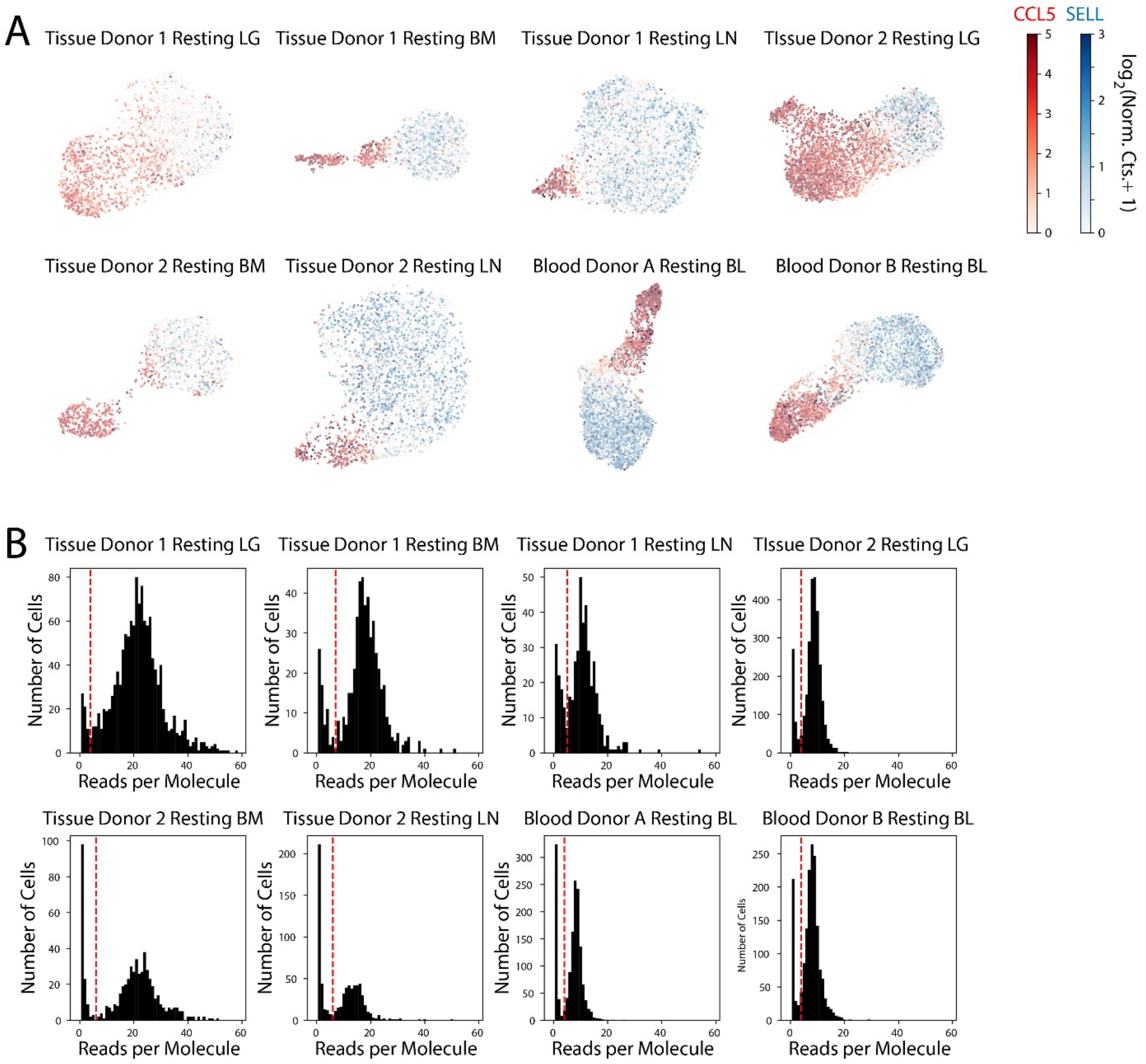


**Supplementary Figure 4. Analysis of CCL5$^+$ effector memory T cells.** (A) UMAP embeddings of the resting T cells from each sample from each tissue and blood donor where each cell is colored by expression of *CCL5* (blue color bar) and *SELL* (red color bar). The two genes, which mark effector memory and non-effector memory populations, respectively, are essentially mutually exclusive. (B) Histograms of the average number of reads per molecule across cells in each sample from each tissue and blood donor for *CCL5*. The distributions are universally bimodal with the lower mode likely arising from molecular cross-talk. The dashed red line indicates the threshold below which the detection of *CCL5* was considered to be artifactual (see Online Methods) for each sample.

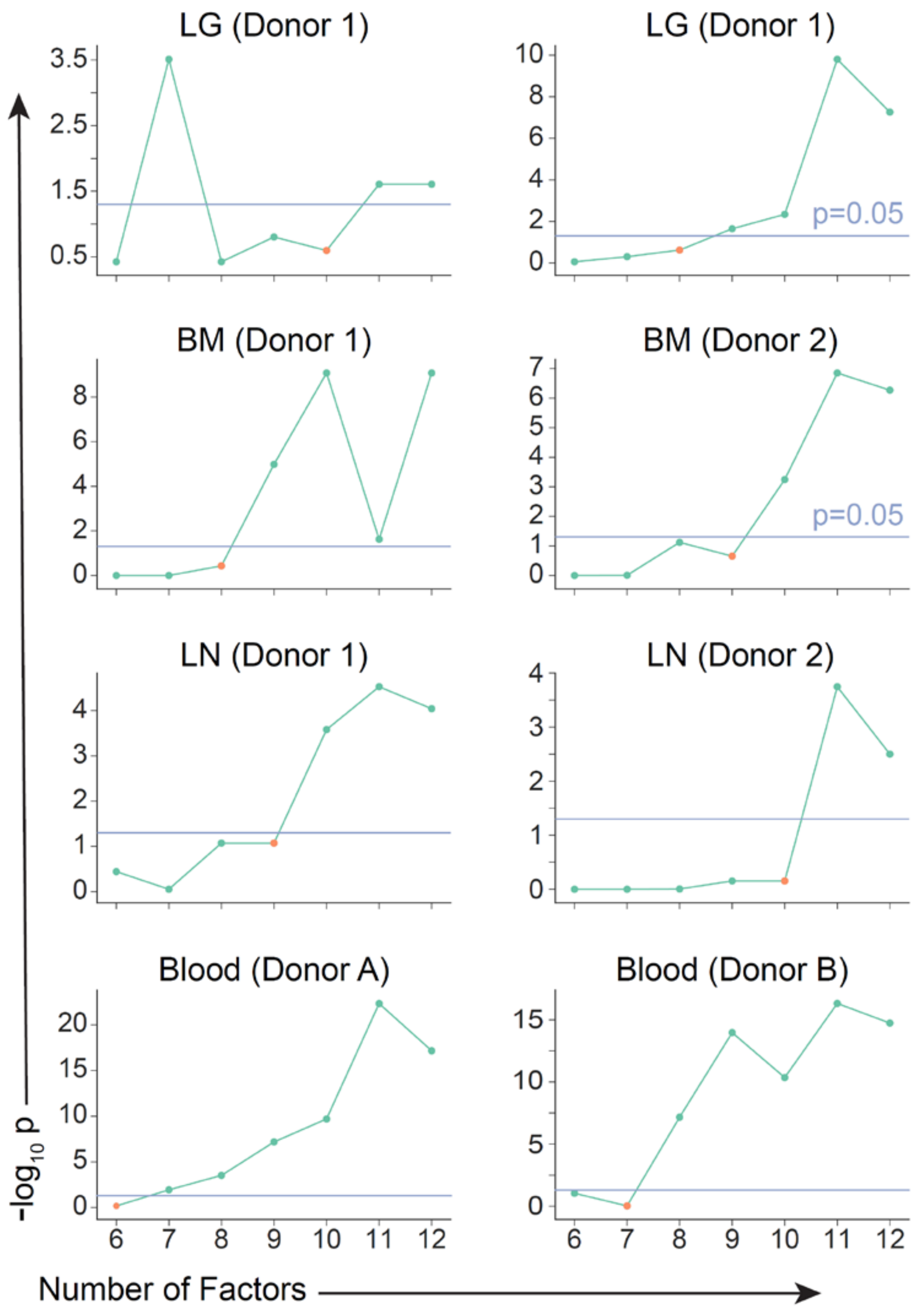


**Supplementary Figure 5. Selection of the number of factors, *K*, for scHPF analysis of each sample.** K was chosen to be the lowest value for which the p-value for pairwise overlap between the top 300 genes in each factor never exceed 0.05 based on the hypergeometric test (see Online Methods). For each sample, the selected value of *K* is indicated by a red dot.

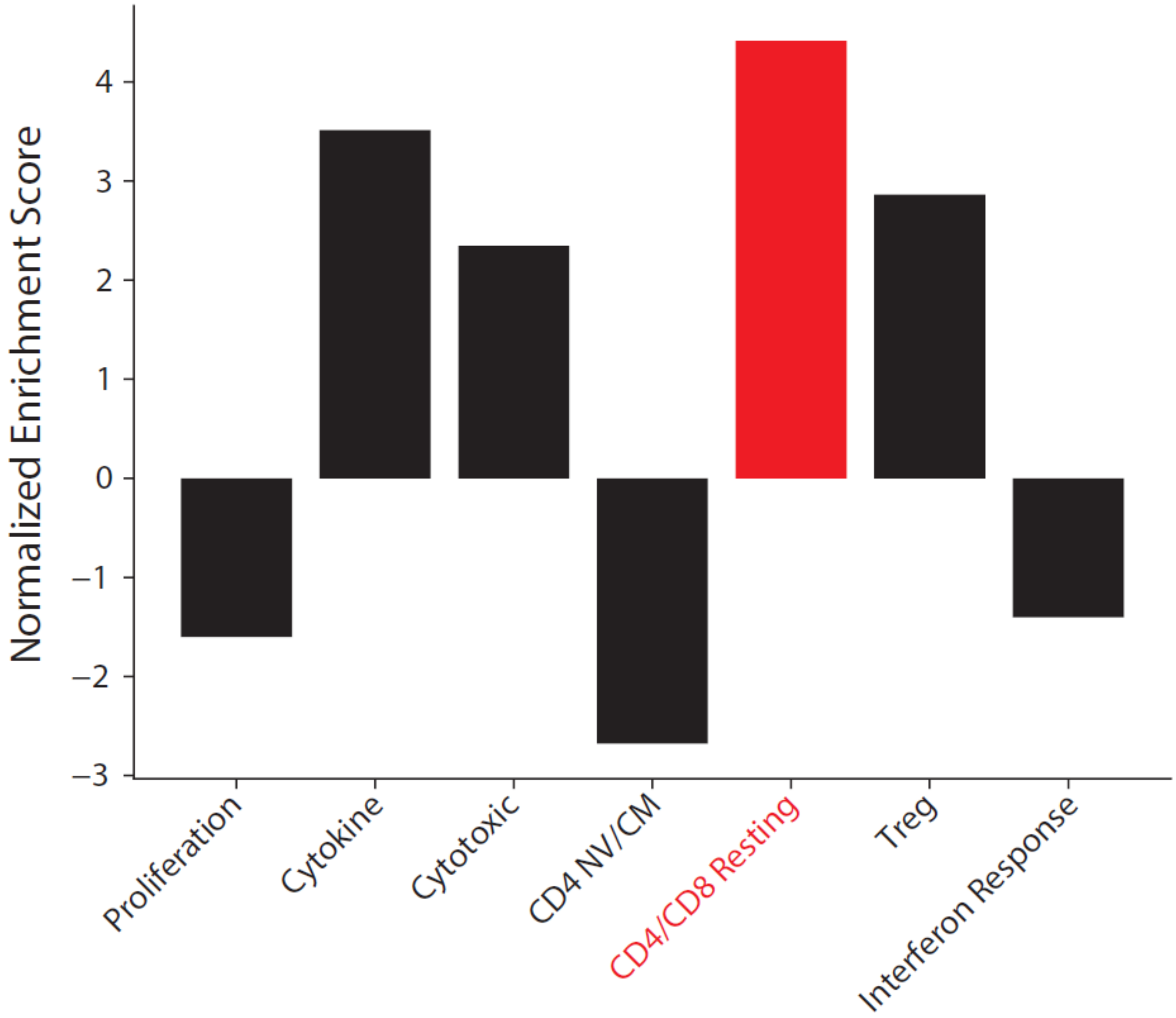


**Supplementary Figure 6. Enrichment of tissue T cell signature in modules.** We used gene set enrichment analysis (GSEA) to assess the enrichment of the tissue T cell signature from Fig. 3 in ranked gene lists for each module (genes ranked by scHPF gene score). The CD4/CD8 Resting module (red), which lacks factor from the blood, exhibited the highest enrichment ($p < 0.00001$) of any module.

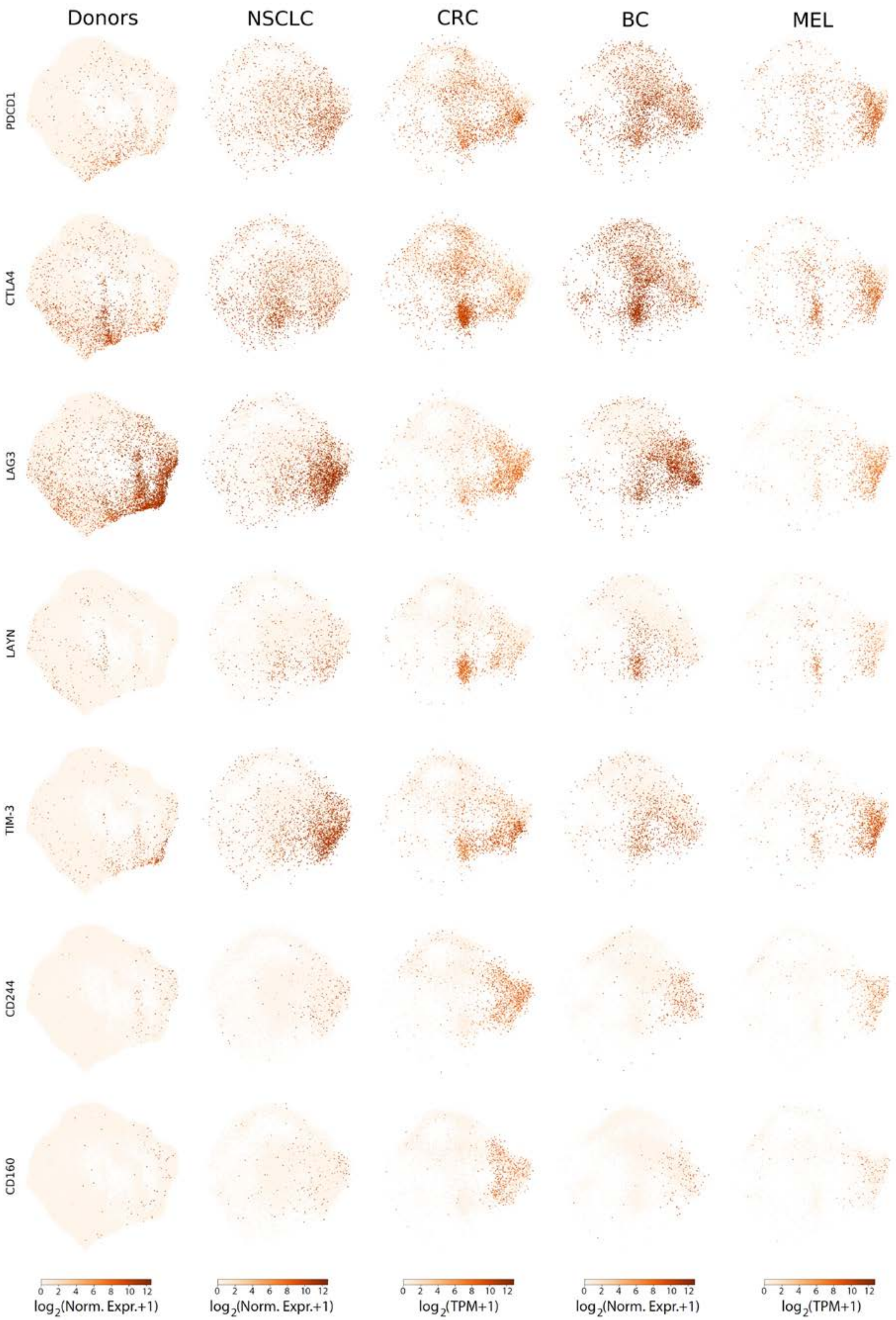
Donors
NSCLC
CRC
BC
MEL
PDCD1
CTLA4
LAG3
LAYN
TIM-3
CD244
CD160
0 2 4 6 8 10 12
log2(Norm. Expr.+1)
log2(Norm. Expr.+1)
log2(TPM+1)
log2(Norm. Expr.+1)
log2(TPM+1)

**Supplementary Figure 7. Expression of exhaustion markers in organ and blood donor T cells and tumor-associated T cells.** Using the UMAP embedding of the merged organ donor and blood T cells from Fig. 6 and the UMAP projections of the tumor-associated T cells from four tumor types onto this embedding, we colored individual cells in each dataset based on expression of T cell exhaustion markers.

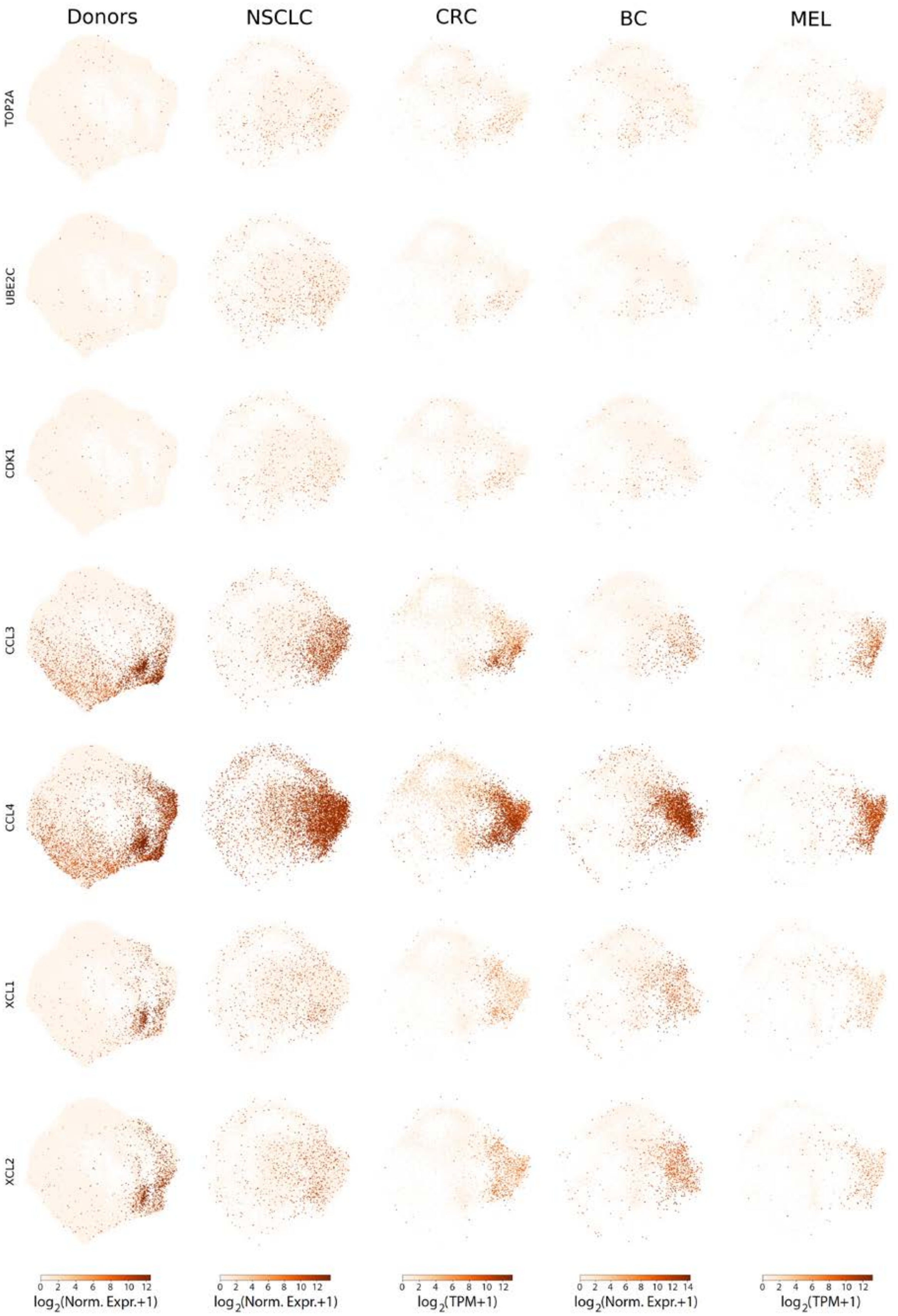

Donors
NSCLC
CRC
BC
MEL
TOP2A
UBE2C
CDK1
CCL3
CCL4
XCL1
XCL2
0 2 4 6 8 10 12
log2(Norm. Expr.+1)
0 2 4 6 8 10 12
log2(Norm. Expr.+1)
0 2 4 6 8 10 12
log2(TPM+1)
0 2 4 6 8 10 12 14
log2(Norm. Expr.+1)
0 2 4 6 8 10 12
log2(TPM+1)

**Supplementary Figure 8. Expression of cell cycle and chemokine genes in organ and blood donor T cells and tumor-associated T cells.** Using the UMAP embedding of the merged organ donor and blood T cells from Fig. 6 and the UMAP projections of the tumor-associated T cells from four tumor types onto this embedding, we colored individual cells in each dataset based on expression of cell cycle markers (*TOP2A*, *UBE2C*, *CDK1*) and chemokines (*CCL3*, *CCL4*, *XCL1*, *XCL2*).

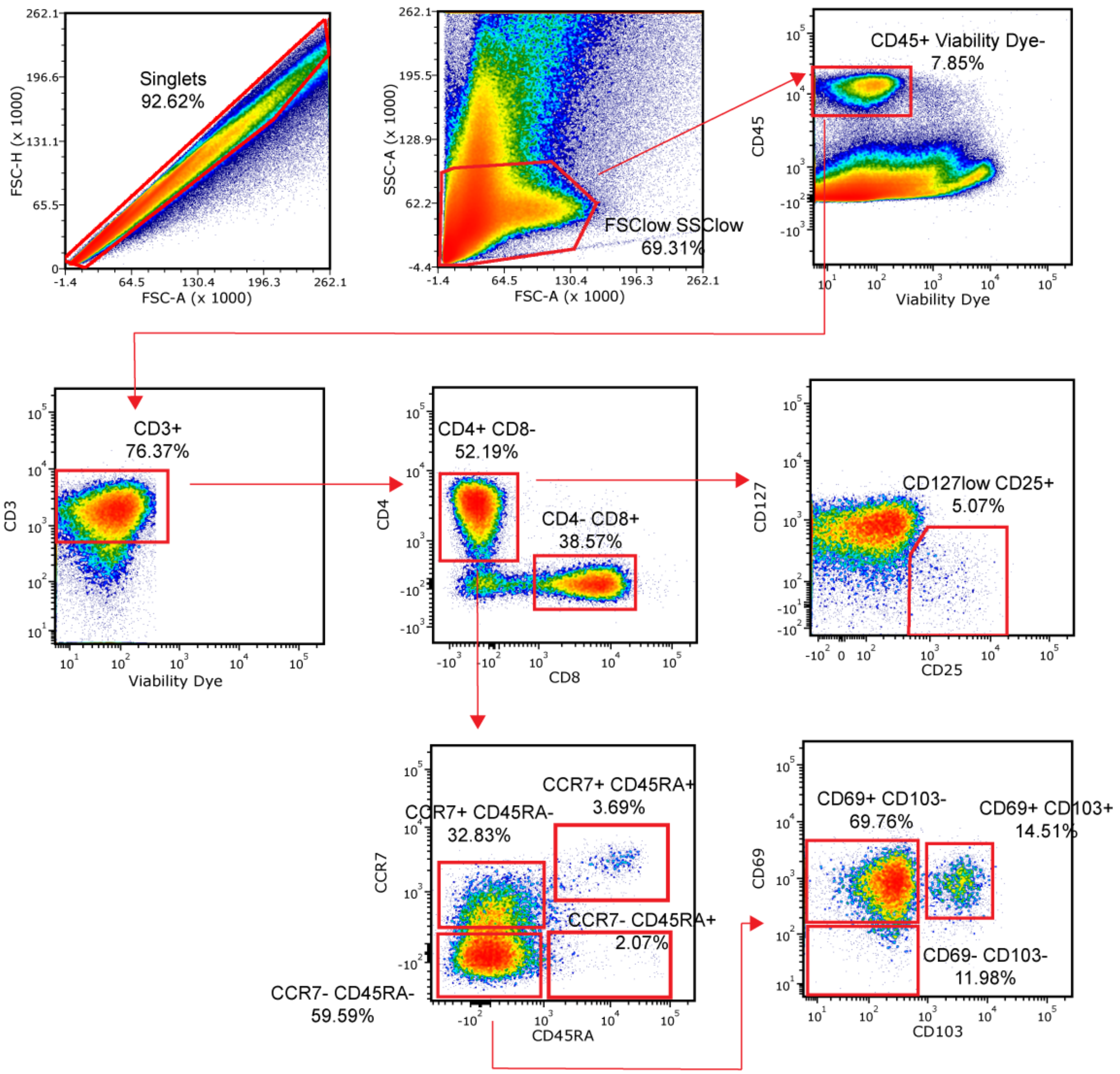


**Supplementary Figure 9. Phenotypic analysis of blood and tissue T cells.** Relative proportions of CD4$^{+}$T cell subsets isolated from the lungs of Tissue Donor 1 shown as an representative gating strategy. All samples were first gated on singlets, FSClow/SSClow, CD45$^{+}$ and Viability Dye$^{-}$ (indicating live cells) before staining for CD3 and subsequent lineage and subset markers. All samples from both Tissue Donors and Blood Donors were analyzed similarly. Data was acquired on a BD LSRII instrument and analyzed by FCS Express software (see Online Methods).

| | Tissue Donor 1 | Tissue Donor 2 |
|---|---|---|
| **Demographics** | | |
| Sex | Male | Male |
| Age (years) | 65 | 52 |
| Ethnicity/race | White | Hispanic/Latino |
| Body Mass Index (kg/m$^2$) | 24.1 | 30.7 |
| **Clinical Characteristics** | | |
| Cause of Death | Cerebrovascular Accident/Stroke | Head Trauma |
| Mechanism of Injury | Intracranial Hemorrhage | Gunshot Wound |
| CPR administered | – | + |
| **Comorbidities** | | |
| Hypertension | + | + |
| Diabetes | – | – |
| CAD | – | – |
| **Social History** | | |
| Smoking History | – | + |
| Alcohol Use | + | + |
| I.V. Drug Use | – | – |
| **Serology** | | |
| CMV | – | + |
| EBV | + | + |
| Toxoplasma | – | – |

CAD, coronary artery disease; CMV, cytomegalovirus; CPR, cardiopulmonary resuscitation; EBV, Epstein Barr virus.

**Supplementary Table 1: Demographic and clinical information from human organ donors.**

| | Tissue Donor 1 | Tissue Donor 2 |
|---|---|---|
| **Lung (Resting/Activated)** | | |
| Total Cells | 3,809 / 2,463 | 5,577 / 4,365 |
| T cells | 2,488 / 1,446 | 4,089 / 3,036 |
| T cell fraction | 0.653 / 0.587 | 0.710 / 0.696 |
| Average Transcript Molecules Detected (T cells) | 2,045 / 3,511 | 2,353 / 3,622 |
| **Bone Marrow (Resting/Activated)** | | |
| Total Cells | 2,251 / 2,512 | 1,918 / 2,191 |
| T cells | 1,826 / 2,080 | 1,304 / 1,455 |
| T cell fraction | 0.811 / 0.828 | 0.680 / 0.664 |
| Average Transcript Molecules Detected (T cells) | 3,666 / 4,359 | 2,272 / 3,690 |
| **Lymph Node (Resting/Activated)** | | |
| Total Cells | 4,896 / 4,649 | 3,395 / 5,820 |
| T cells | 4,194 / 4,186 | 2,992 / 5,190 |
| T cell fraction | 0.857 / 0.900 | 0.881 / 0.892 |
| Average Transcript Molecules Detected (T cells) | 3,126 / 2,888 | 3,320 / 3,789 |

| | Blood Donor A | Blood Donor B |
|---|---|---|
| **Blood (Resting/Activated)** | | |
| Total Cells | 4,872 / 5,411 | 4,777 / 4,775 |
| T cells | 4,282 / 4,911 | 4,196 / 4,236 |
| T cell fraction | 0.879 / 0.908 | 0.878 / 0.887 |
| Average Transcript Molecules Detected (T cells) | 4,131 / 6,082 | 4,256 / 6,342 |

**Supplementary Table 2: scRNA-seq cell numbers, T cell purity, and transcripts detected.**
